# The key to complexity in interacting systems with multiple strains

**DOI:** 10.1101/2020.02.03.932806

**Authors:** Erida Gjini, Sten Madec

## Abstract

Ecological community structure, persistence and stability are shaped by multiple forces, acting on multiple scales. These include patterns of resource use and limitation, spatial heterogeneities, drift and migration. Pathogen strains co-circulating in a host population are a special type of an ecological community. They compete for colonization of susceptible hosts, and sometimes interact via altered susceptibilities to co-colonization. Diversity in such pairwise interaction traits enables the multiple strains to create dynamically their niches for growth and persistence, and ‘engineer’ their common environment. How such a network of interactions with others mediates collective coexistence remains puzzling analytically and computationally difficult to simulate. Furthermore, the gradients modulating stability-complexity regimes in such multi-player systems remain poorly understood. In a recent study, we presented an analytic framework for *N*-type coexistence in an SIS epidemiological system with co-colonization interactions. The multi-strain complexity was reduced from *O*(*N*^2^) dimensions of population structure to only *N* equations for strain frequency evolution on a long timescale. Here, we examine the key drivers of coexistence regimes in such a system. We find the ratio of single to co-colonization *μ* critically determines the type of equilibrium for multi-strain dynamics. This key quantity in the model encodes a trade-off between overall transmission intensity *R*_0_ and mean interaction coefficient in strain space *k*. Preserving a given coexistence regime, under fixed trait variation, can only be achieved from a balance between higher competition in favourable environments, and higher cooperation in harsher environments, consistent with the stress gradient hypothesis in ecology. Multi-strain coexistence regimes are more stable when *μ* is small, whereas as *μ* increases, dynamics tends to increase in complexity. There is an intermediate ratio that maximizes the existence and stability of a unique coexistence equilibrium between strains. This framework provides a foundation for linking invariant principles in collective coexistence across biological systems, and for understanding critical shifts in community dynamics, driven by simple and random pairwise interactions but potentiated by mean-field and environmental gradients.

## Introduction

Rich ecosystems comprise many species interacting together in a myriad of ways and on multiple temporal and spatial scales. Understanding the scope and consequences of such interactions has been the focus of countless theoretical ecology studies, starting with the seminal work by Lotka (Lotka, 1926) and Volterra (Volterra, 1926) on mathematical models of the population dynamics of interacting species. This model has been later extended and sophisticated by many other theoretical studies (May, 1972; Pascual et al., 2006), and is currently extensively used to characterize interaction networks in empirical microbiome communities (Bucci et al., 2016; Stein et al., 2013). Theoretically, a crucial question has been to study stability and coexistence patterns in such Lotka-Volterra multi-species communities, analyzing both structured ecological networks and random networks (Serván et al., 2018). Modeling efforts seek to understand organizing principles for species composition, such as the balance between competition and cooperation (Mougi and Kondoh, 2012), which many are increasingly arguing that should be studied together in integrated and context-dependent frameworks (Bascompte, 2019; Coyte and Rakoff-Nahoum, 2019). Overall, analysis of such models with arbitrarily high dimensionality has been and continues to remain difficult.

The challenge of high-dimensionality in ecological microbial networks parallels a similar challenge in the epidemiology of polymorphic pathogen systems, where understanding the mechanisms and forces that maintain diversity among interacting strains, is also an area of active research (Cobey and Lipsitch, 2012; Gupta and Anderson, 1999; Lipsitch et al., 2009; Wearing and Rohani, 2006). It is well recognised that population patterns of infection are to a large extent determined by susceptibility to infection. Most multi-strain SIR compartmental models, inspired for example from influenza, dengue, and malaria parasites, have focused on cross-immunity thresholds and thus competition between antigenically similar strains as the main driver of strain interactions and pathogen population structure (Gupta et al., 1998). To deal with the inherent mathematical complexity, such studies have adopted model reduction, traveling wave solutions (Gog and Grenfell, 2002), and aggregation of variables (Gomes et al., 2002; Lin et al., 1999), which have led to key insights and analytical results. Studies have shown that even in a simple transmission model with a single infectious agent, where immunity by infection and vaccination reduce susceptibility to subsequent infections, altered susceptibility induces a ‘reinfection threshold’, above which reinfection dominates transmission, leading to higher infection levels and vaccination failure (Gomes et al., 2004). It is unsurprising that if multiple strains are involved, very complex behavior can emerge in infectious disease systems when such nonlinearities multiply and interact. Yet, it remains unclear which other factors, besides persistent host immunity, make strains compete or cooperate with each other, and whether there are environmental factors that can shift their fitness balance epidemiologically.

Here, we set out to study another mode of strain interactions, that has received, albeit less, modeling attention, namely interactions between strains via altered susceptibilities, not after, but during infection, in endemic scenarios of no immunity (SIS). Co-colonization vulnerability in SIS models with up to 2 strains, has been shown to be important in several studies (Alizon et al., 2013; Davies et al., 2019; Gaivão et al., 2017; Gjini et al., 2016; Lipsitch, 1997), but very few analytical investigations have been done for a larger number of interacting strains (Adler and Brunet, 1991). In a recent mathematical study, extending the cocolonization interaction space to *N*-strains, we have provided a new analytical front for understanding a complex SIS system of multiple strains interacting upon co-colonization (Madec and Gjini, 2019). Using timescale separation, we obtained a model reduction, where we showed how from the matrix of multi-type interactions arising in SIS cocolonization, we can obtain a special replicator equation (Cressman and Tao, 2014; Hofbauer and Sigmund, 2003) for strain frequencies. This reduced model formulation makes the entire dynamics more accessible to analysis, and links directly epidemiological SIS dynamics to Lotka-Volterra systems (Bomze, 1995) and evolutionary dynamics (Nowak and Sigmund, 2004).

In the present article, we harness the simplicity of this co-colonization model framework to investigate coexistence and stability of such multi-strain systems, without immunity, but with variable co-colonization susceptibility coefficients among strains. We start by studying the behavior of the system for different global environmental variables such as total transmission intensity *R*_0_, and mean interaction coefficient in the pool of available strains k. We then study coexistence through random co-colonization interactions, where the matrix coefficients are drawn from fixed distributions, and can range from competitive to cooperative links. We ask what is the number of strains that can coexist when starting from a pool of *N* strains, and in which diversity-stability configuration. We uncover rich transient and asymptotic behaviour of such systems, where steady states, limit cycles, multi-stability and chaotic attractors are possible.

We argue that the ratio of single-to co-colonization is a critical factor in collective dynamics, by modulating the asymmetry in pairwise mutual invasibility outcomes between types, and consequently, the dynamic complexity of the system as a whole. We argue that the analytically explicit form of this ratio in our model enables direct connection with the stress gradient hypothesis (SGH) in ecology (Bertness and Callaway, 1994; Callaway and Walker, 1997). This hypothesis postulates that as stress increases, the importance of positive facilitative effects increases in a community, whereas in benign environmental conditions, competitive effects are higher; a finding that emerges also from our results. In support of complex higher-order dynamics emergent from simple pairwise interactions, with critical links between mean and variance, we uncover the exact formulation for why the sum as a collective is much more than its parts. Our results invite a deeper understanding of the biology of such multi-strain systems and point to key global modulators of collective polymorphic coexistence in nature.

## Results

In the present model, the entire system is structured as a collection of hosts that can be in different colonization states: susceptible, singly-colonized or co-colonized. We consider a multi-type infection, transmitted via direct contact, following susceptible-infected-susceptible (SIS) epidemiological dynamics with co-colonization (Madec and Gjini, 2019). Adopting a general formulation for diversity, we assume there are *N* types, without specifying the mechanisms for their definition. Thus, with an ordinary differential equations model (see Materials and Methods), we describe the proportion of hosts in several compartments: susceptibles, *S*, hosts colonized by one type *I_i_*, and co-colonized hosts *I_ij_*, with two types of each combination, independent of the order of their acquisition. Fitness differences between types are encoded in how strains interact with each other upon cocolonization (*K_ij_*), - whether there is facilitation between resident and co-colonizer (*K_ij_* > 1) or inhibition (*K_ij_* < 1), - assuming equivalence in transmission *β* and clearance rate *γ*. The structure of the *K_ij_* is central to multi-strain dynamics. The co-colonization interaction matrix between *N* strains in the system can arbitrarily include cooperative and competitive entries, but subject to small deviation from neutrality *K_ij_* = *k* + *εα_ij_* (*ε* small). We take no account of other biological details such as mutation, seasonality in transmission, or heterogeneity in the host population, which may influence the transmission dynamics of particular strains. These exclusions serve our purpose to assess the impact of selection imposed by co-colonization interactions between strains in the host population on temporal trends in individual strain frequencies.

In an earlier mathematical investigation (Madec and Gjini, 2019), we have derived in detail two timescales in this system: a fast one, given by the neutral model, and a slow one governed by the variation in co-colonization coefficients *K_ij_*. During fast dynamics the system stabilizes the total prevalence of single and co-colonization at the endemic persistence equilibrium, whereas over the slow timescale, under conserved global quantities, strain selective dynamics unfold. In the *N* equations, describing frequency evolution, the changing mean fitness of the collective appears explicitly, highlighting a key environmental feedback on all strains (see Box 1).

### Box 1. Key model features for *N*-strain cocolonization and dynamics

- **Deviation from neutrality in interaction trait space** The basis of the framework (Madec and Gjini, 2019) is to conceptualize each interaction coefficient between closely-related strains in co-colonization, as a mean value *k* plus some deviation from neutrality, thus re-writing it as:

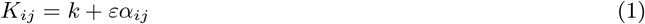

where 0 < *ε* < 1 is small. Thus, *A* = (*α_ij_*) is the normalized interaction matrix, relative to the reference *k*. The parameter *k* is an indicator of mean interaction in co-colonization between any two strains, which when above 1 describes facilitation, and when below 1 describes competition. Another key epidemiological parameter in the model is the basic reproductive number, *R*_0_, describing the transmissibility of infection, defined as the average number of secondary infections that arise from single case in a naive population (Diekmann et al., 1990).
- **Fast and slow dynamics: derivation of the replicator equation in a new context** On the fast time-scale (*ε* = 0), the system tends towards neutrality and reaches the manifold where total prevalence of susceptibles **S**, total prevalence of single colonization **I** and total prevalence of co-colonization **D** are conserved, and depend only on mean epidemiological parameters:

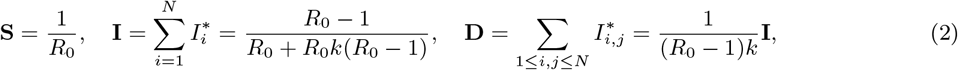

but, where each individual strain frequency is neutrally stable and free to vary *z_i_* ∈ [0,1]. On a second time scale, which is the slow time scale *τ* = *εt*, consistent with weak selection, exact asymmetries between strains play out, and while the conservation law above still holds, individual strain frequencies z¿ are not free anymore. They obey deterministic slow dynamics, given the *N*-dimensional system:

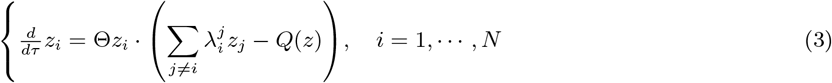

where 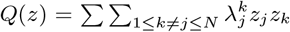, is a quadratic term symmetric on all strains, encapsulating the effect of the system as a whole on each individual strain. We have 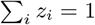 and the rate Θ is given by: 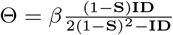.
- **From dyads to collective dynamics** In the above equations, which drastically reduce the system from *N*(*N* – 1)/2 + *N* to only *N* dimensions, the slow dynamics of strain frequencies are a direct function of pairwise invasion fitnesses 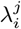 between strains, which in our model, for each dyad (*i,j*), have the simple form:

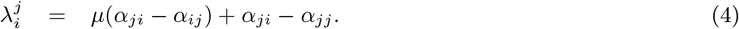 The quantities 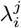 denote the initial rate of growth of strain *i* in an exclusion equilibrium where only strain *j* is resident, and depend on the ratio between single and co-colonization *μ* which is given by

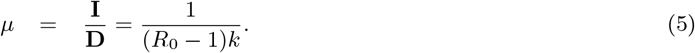
- **Invariant principles in non-equilibrium multi-strain dynamics** On the slow timescale, at all times, and for all strains, the following relationships hold:

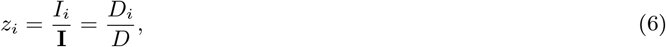

stating that *z_i_*, as a measure of dominance of strain *i* in the total prevalence of carriage, occupies an equal relative frequency in single (*I_i_*) and co-colonization 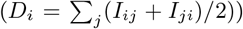. After solving for strain frequencies *z_i_*, the Eqs. 3 can inform the epidemiological variables as follows:

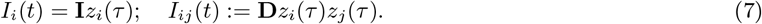

### The influence of mean-field parameters on tempo and type of coexistence

According to this model, global mean-field parameters, such as those affecting transmission rate, *β*, basic reproduction number *R*_0_, and mean interaction coefficient *k* affect explicitly the multi-strain dynamics over long time scales (*εt*). The equation for strain frequencies *z_i_* contains information for how they set the speed (Θ) and mode of strain stabilization (*μ*). More specifically, the relative dominance of single colonization in the system *μ*, because it appears as a factor in pairwise invasion fitness 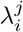, modulates the importance of cross-strain asymmetry in co-colonization trait comparison between two strains: the higher *μ* is, the higher the dominance of single colonization, the more important the asymmetry between *i* and *j* becomes (*a_ji_* – *a_ij_*). While this first component of invasion fitness is sensitive to feedbacks from global transmission and mean parameters between strains, the other component of the 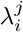 is independent of such global quantities and depends only on characteristics of the resident, namely on the relative strength of inter-vs. intra-strain interaction of the resident (*a_ji_* – *a_ij_*).

Notice, that for fixed normalized variation among strains, hence rescaled matrix *A* = (*a_ij_*), increasing *μ* in the system, amplifies asymmetries and should lead to more competitive exclusion between strains. We illustrate this for the case of *N* = 2 in Fig. 1a, where for randomly generated rescaled interaction matrices *A* (Normal distribution with mean 0 and variance *ε*), increasing *μ* stretches the region occupied by 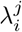 towards the competitive exclusion zone (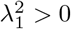, and 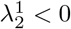). This phenomenon escalates also to the case of higher *N*, where effectively the multi-strain network is derived from pairwise invasion fitnesses between any two constituent strains (Fig. 1b). As the ratio of single to co-colonization, *μ*, increases, the proportion of edges between constituent strains, that lead to competitive exclusion increases, making collective coexistence subject to more constraints.

**Figure 1:**
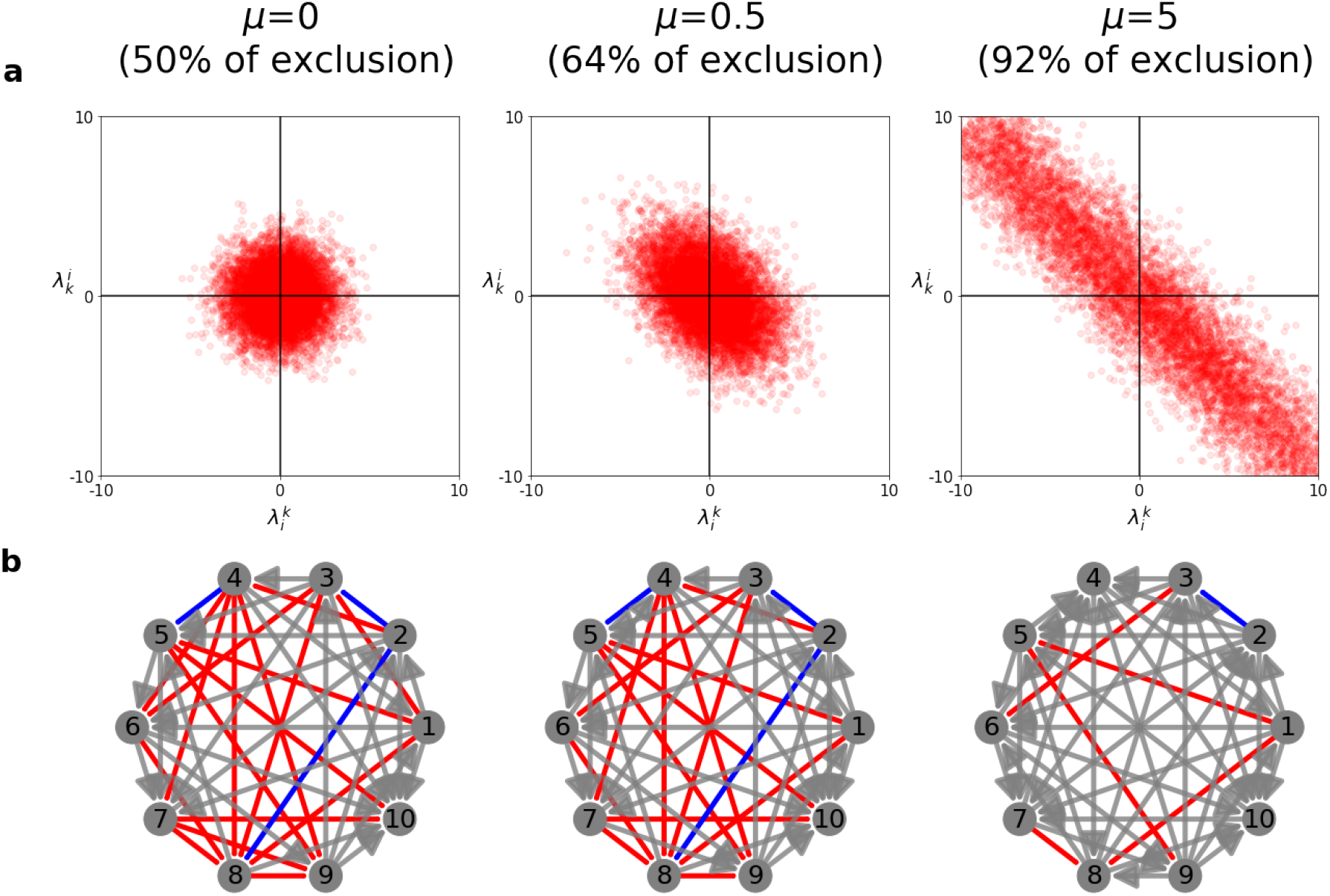
Collective coexistence from pairwise invasion fitnesses and the effect of single-to cocolonization ratio *μ*. **a.** Effects of *μ* on the repartition of the pairwise interactions. In our model, there are four possible outcomes between two strains (depending on the values of 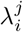): extinction of 1, extinction of 2, coexistence or bistability. The coefficients *α_ij_* of the matrix *A* = (*α_ij_*) are generated following a normal distribution 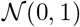. When *μ* → 0, the partitioning between the four outcomes is the same. When *μ* increases, the strain pairs, for the same *A* matrix, follow more likely competitive exclusion dynamics. **b.** Effect of *μ* on the multi-strain pairwise λ networks. Red line: coexistence, blue line: bistability, and a grey arrow is a competitive exclusion (the arrows shows the winner between the two). Here we fixed the normalized interaction coefficients between strains (matrix *A*) and *N* =10 strains and we varied *μ*. As *μ* increases even though the actual normalized interaction coefficients remain fixed, the effective outcomes between each pair of strains, being *μ*-dependent, tend to exclusion, with the grey edges becoming more common. This feature makes stable coexistence harder for *μ* large.

Although the mean number of strains that can coexist (*n*) apparently increases with *μ* (See Supplements for *N* = 10), the possibilities for stable coexistence tend to be more and more restricted to only special numbers of strains (odd values), as expected generically in special cases of the replicator equation. To illustrate further the role of *μ*, we show the dynamics of strain frequencies as a function of *μ* in Fig. 2, for the case of *N* = 3 and *N* = 4. Under fixed asymmetries *A*, changing the ratio between single and co-colonization, shifts the multi-strain dynamics from stable coexistence towards limit cycles and ultimately towards May-Leonard type oscillations, with effectively only one strain persisting for long periods of time, to be subsequently replaced by another one, and so on. These extreme regimes of behavior are expected in the replicator equation for special cases, but here the novelty is that we have identified a global environmental quantity, the ratio between single and co-colonization, as a tuning parameter for moving our epidemiological system between such extremes (see Box 2).

**Figure 2:**
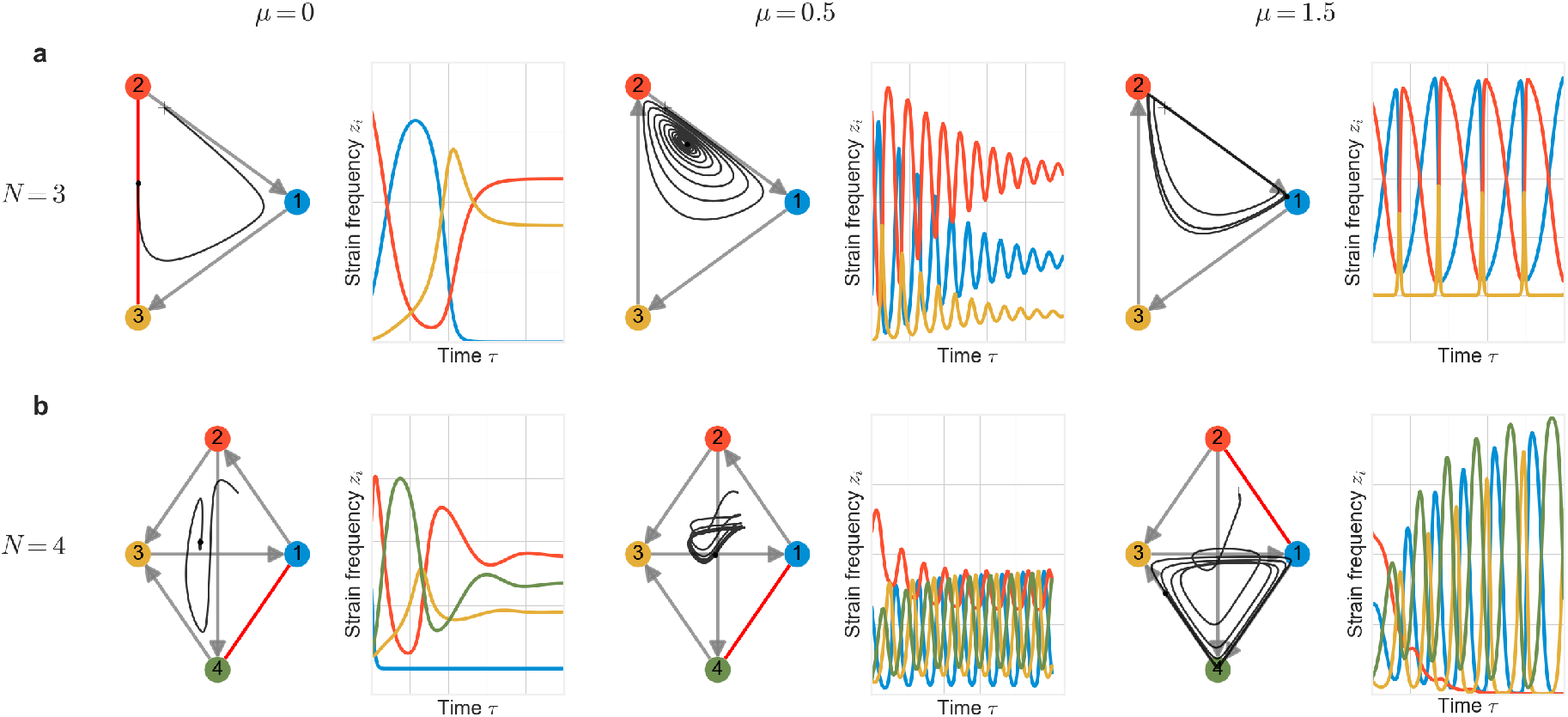
Increasing the ratio of single to co-colonization (*μ*) increases the complexity of multi-strain dynamics. We illustrate two examples of fixed interaction matrix *A* and shifting *μ*. **a.** for *N* = 3 and **b.** for *N* = 4. Here we fixed the normalized interaction coefficients between strains (matrix *A*) and we varied *μ*. Because *μ* affects the 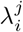, for each *μ* we have a different effective competition network between strains modulated by top-down factors. The plots show the strain frequency (*z_i_*) dynamics following the equations on the slow manifold. For *N* = 3 and *N* = 4, for small values of *μ*, the final attractor is a stable steady state, while for a large value of *μ*, the stable attractor is a union of heteroclines of 3 strains (May Leonard type). For intermediate values of *μ* (*μ* = 0.5) we obtain a stable spiral of coexistence for *N* = 3 and a limit cycle of coexistence for *N* = 4, both consistent with an increase in rate of strain turnover.

#### Box 2. Single to co-colonization ratio as a tuning parameter for complexity

One limit for *μ* may be achieved by either increasing cooperation *k* → ∞, or increasing transmission intensity 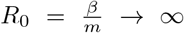. The other limit can be achieved by increasing competition or decreasing transmission intensity. Although in reality, these limits may never be literally attained biologically, here we describe the mathematical trends of the system, which will most likely reside in the intermediate range. For a more detailed analysis of system behavior and speed of the dynamics Θ_*lim*_ in these limits, see also Supplementary Text S2, and Figures S2–S3. A specific example with dynamics close to these limits is provided in Figure S4.

- **Limit 1: Co-colonization,** *μ* → 0 (*R*_o_ ↑ **or** *k* ↑) <u>Quality of the dynamics</u> Recall that 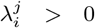 implies that the strain *i* (the invader) may invade the system with only strain *j* (the resident) present, starting from a very small frequency. In the limit *μ* → 0, this pairwise invasion fitness reduces to:

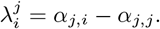 Importantly, in this limit, the fitness 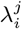 depends only on the coefficients of the resident, where *α_j,i_* measures how much the resident *j* contributes to mixed cocolonization with strains *i* and *j*: *D_ij_* (transmission and fitness of other strains), and *α_j,j_* how much the resident *j* contributes to self co-colonization: *D_jj_* (transmission and fitness of itself). Hence, the relative fitness of the invader *i* depends only on the resident *j*. Invasion is possible if and only if *α_j,i_* > *α_j,j_*. This phenomenon in a system of *N* players can increase the availability of niches. For example, if all intra-specific coefficients are higher than inter-specific coefficients: *α_j,j_* > *α_i,j_* for all *j* = 1, ··· *N* and *i* ≠ *j* then all strains may be stable residents when alone, and we would have at least *N* stable monomorphic equilibria. Cycles or more complex dynamics are rare. The final outcomes can vary because the structure of the matrix Λ is free in principle. In the special case *α_jj_* = 0 for all *j*, then Λ = *A^T^*, and the dynamics could be richer, in line with generic replicator systems (Yoshino et al., 2008). In general, for random A, with no special structure, qualitatively, the system likely has many stable steady states wherein only few species coexist. <u>Speed of the dynamics</u>

i. Θ_*lim*_ = 0. *If k* → ∞, *we have* 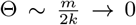. *For fixed population turnover (m: clearance and death rate), the selective dynamics happen very slowly, i.e. the dynamics are effectively neutral. Similarly if R*_0_ → ∞ *via decreases in m (m* → 0), *the multi-strain system stays very close to neutrality*.
ii. Θ_lim_ > 0. *If R*_0_ → ∞, *via increases in transmission rate, β* → ∞, *then* 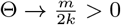 *which is far from zero. In this case, the model reduction captures well the real dynamics and the qualitative study of μ* → 0 *is appropriate*.
- **Limit 2: Single colonization,** *μ* → ∞ (*R*_0_ ↓ **or** *k* ↓) <u>Quality of the dynamics</u> Recall the matrix Λ is the matrix of all pairwise invasion fitnesses between *N* strains in the system. In this limit, we have generically ║Λ║ → +∞ and Θ → 0. In order to keep a bounded matrix Λ we rewrite the system as

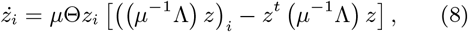

where *i* = 1, ···, *N*. Then the speed is given by *μ*Θ and the qualitative behaviour by *μ*^-1^Λ. Notice that when *μ* → +∞, the matrix *μ*^-1^Λ → *A^T^* – *A*. Importantly, this matrix is skew symmetric, for which there are known mathematical results of the replicator equation in zero-sum games (see (Allesina and Levine, 2011; Chawanya and Tokita, 2002; Fisher and Reeves, 1995). The quadratic term in Eq. 8 is zero: *Q*(*z*) = *z^T^*Λ*z* = 0. So the *N*-strain frequency dynamics reduce to:

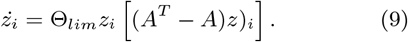 There exists exactly one nonnegative linearly stable equilibrium (in particular multistability is impossible). In practice, similar to the classical Lotka-Volterra model, there is a one-parameter family of limit cycles parametrized by initial conditions. This type of coexistence is not structurally stable, and will be lost for a large but finite value of *μ* leading in general to heteroclinic limit cycles among strains. Denoting *n* the number of strains coexisting at equilibrium, for this skew-symmetric case of the replicator equation, it has been shown that only an odd number of strains may coexist (see also Fig. S1, S5). More specifically, the probability to observe *n* = *k* strains, out of a total pool of *N* is

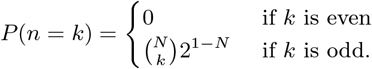 <u>Speed of the dynamics</u>

i. Θ_*lim*_ =0. *If R*_0_ → 1 *decreases transmission intensity, either via lower β or higher m, then* Θ → 0, *and selective dynamics are too slow, making the system effectively neutral*.
ii. Θ_*lim*_ > 0 *If k* → 0, 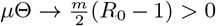. *Thus the selective dynamics are well defined and occur on a feasible timescale*.

Thanks to the explicit formula for this critical ratio, we can see that *μ* = **I**/**D** = 1/*k*(*R*_0_ – 1) can change in two ways: i) either by changing basic reproduction number *R*_0_, or by changing *k*, the mean interaction coefficient between strains in co-colonization. If *R*_0_ increases *μ* decreases, and similarly, if *k* increases (strains tending towards facilitation) *μ* also decreases. From the expression for 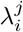, and the illustrations above, *μ* determines completely the type of equilibrium, for a given set of standardized variation in strain interactions. However if *μ* changes, either via shifts in *R*_0_ or in *k*, the speed of the actual selective dynamics between strains may also change. In the case of *R*_0_ effects on *μ*, this depends on how *R*_0_ changes, via transmission rate or duration of infection changes. Thus we have 3 cases, whereby changes in *μ* can lead to changes in 0 (See Supplementary Figure 2). When changes in *μ* are mediated by changes in *k*, for high values of *μ*, the speed of selection goes down. When changes in *μ* are mediated by *R*_0_, for *μ* small (i.e. high *R*_0_) there may be a small region where increasing *μ* (either via decreasing *β* or increasing *m*) can actually speed up the dynamics. This then would imply, that for relatively high *R*_0_, reduction in *R*_0_ (via faster population turnover, shorter duration of carriage, or lower transmission rate) not only acts to increase the complexity of the dynamics, but also speed up of the multi-strain selection, thus contributing to quicker settlement of the competitive hierarchies between entities at an equilibrium.

### Coexistence, stability and diversity

The community dynamics resulting from co-colonization interactions between strains can be very complex, and ranges from simple coexistence equilibria to limit cycles and even possible wildly oscillatory dynamics. While there is an ongoing debate in ecology about the relationship between stability and diversity in ecosystems (May, 1972; McCann, 2000; Odum and Barrett, 1971; Tilman and Downing, 1994), here we set out to examine this relationship in our system, assuming random interaction structure between strains, namely the matrix *A* and fixing the total pool size of strains *N*. Thus, we study the model’s rich behaviour for a larger pool of strains in the system, namely *N* =10 (Figure 3). We sample the normalized interaction matrix *A* randomly from a normal distribution. For each interaction matrix, we compute numerically the feasible equilibria of the system (*z_i_* ≥ 0, ∀*i*), and for each equilibrium found, we evaluate its local stability and its associated Shannon entropy 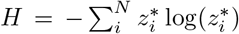. By simulating different randomly generated structures of the matrix, we can explore regimes of exclusion, multi-stability, multi-strain coexistence, limit cycles and even chaos. In this analysis we use the definition of equilibrium stability (McCann, 2000), although there are also other notions of general stability related to permanence (Law and Morton, 1996) that could be informative.

**Figure 3:**
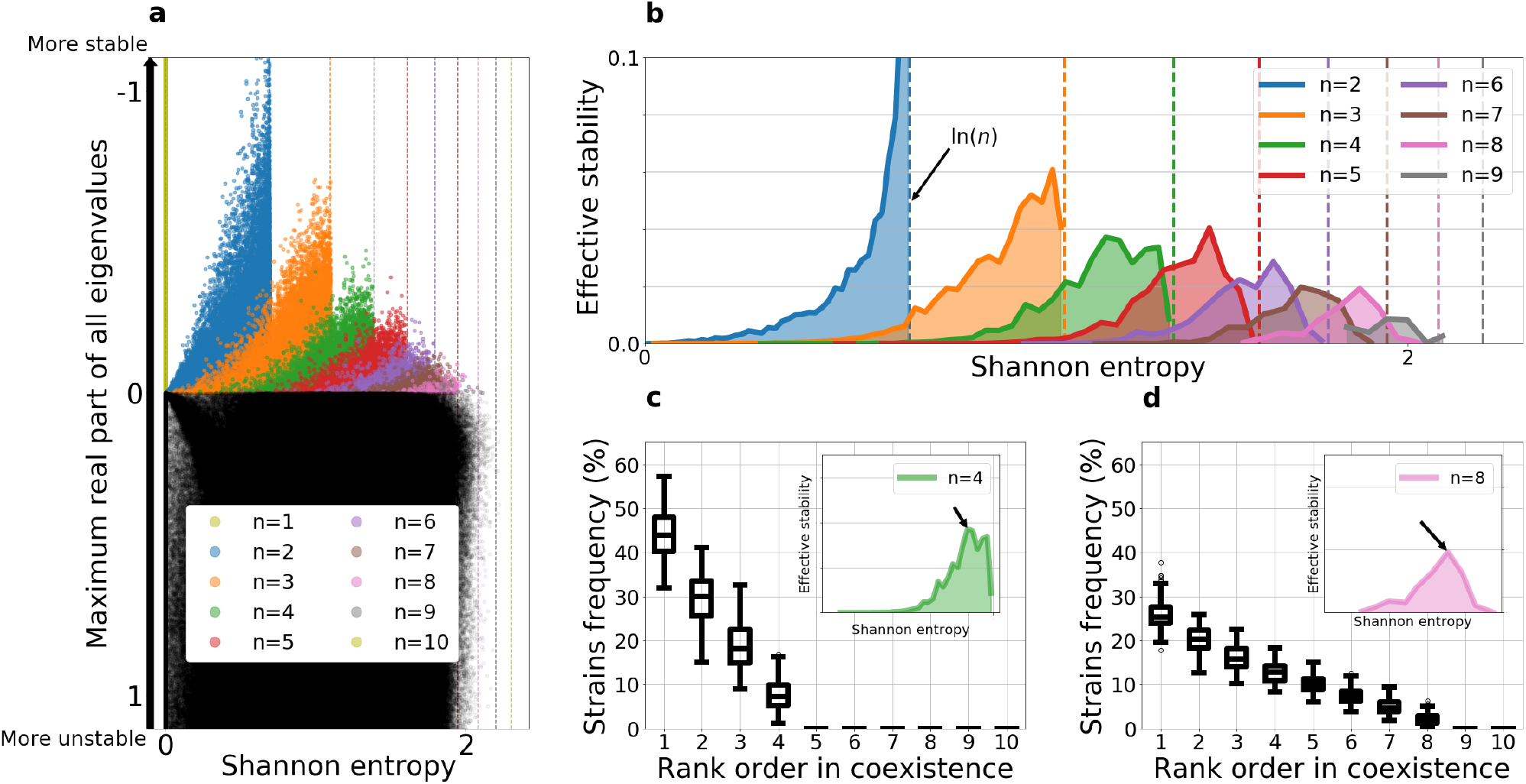
Diversity-stability relationships for *n*-strain coexistence starting with a pool of *N* = 10 strains. **a.** Summary of 100.000 simulations with a random normalized interaction matrix *A*, with each entry *α_ij_* drawn from the Normal distribution 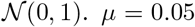. The equilibria of each system are mapped as dots in this plot, after computing their local stability and Shannon entropy. **b.** Effective stability is calculated as a product between the mean dominant eigenvalue and proportion of steady states at each entropy level, stratified by number of strains coexisting *n*. **c.** Plot of the rank-order frequency distribution of strains at the “optimal diversity” for *n* = 4 strains coexisting out of a pool of 10. **d.** Plot of the rank-order frequency distribution of strains at the “optimal diversity” for *n* = 8 strains coexisting out of a pool of 10.

The first aspect of coexistence we focus on is the probability of reaching a stable *n*-strain coexistence, starting with a pool of *N* =10 strains. Using our model, under random allocation of co-colonization interaction coefficients, we find that the probability of stable *n*-strain coexistence decreases with *n*, thus confirming that the more strains there are, with random mutual interactions in co-colonization, the harder it is for all of them to coexist (colored dots in Fig. 3a). However, notice that in contrast to the stable steady states, there are many more unstable steady states in the system (98%), spread at any diversity and stability level, with no apparent order or hierarchy (black dots in Fig. 3a). This suggests that unstable coexistence is much more likely and that the number of coexisting strains in that case is not a limiting factor. For a fixed number of strains coexisting *n*, out of a total pool of *N*, the higher the diversity (Shannon entropy) of the equilibrium, the higher the stability, as measured by the negative part of the dominant eigenvalue of the Jacobian matrix evaluated at that equilibrium.

Then, to quantify in more detail this relationship, accounting for the feasibility of the steady state itself, we go on to examine how the probability of stable coexistence for a given *n* depends on the diversity of the equilibrium. Using Shannon entropy as a measure of diversity of the coexistence equilibrium, we find that among stable equilibria, the proportion of systems at a given entropy level decreases with entropy, and this pattern is consistent for all *n*. This implies that under randomly sampled interactions, stable coexistence equilibria with equally-distributed strain frequencies, and thus more diversity, are harder to obtain.

Thus, on one hand we find more diversity is associated with more stability, in support of other ecological studies (McCann, 2000), but also that higher-diversity equilibria are less feasible and harder to reach under random sampling of interaction coefficients from a pool of *N* strains. When combining these two forces: diverse coexistence is rarer to obtain, but generally more stable, we find that this trade-off between probability of stable coexistence and stability, give rise to an optimal intermediate diversity for *n*-strain coexistence (Fig. 3b)), where stable coexistence is on one hand sufficiently feasible, and the resilience of that equilibrium to perturbations is sufficiently high. Thus, mean stability of *n*-strain coexistence is effectively maximized at particular, typically intermediate, values of Shannon entropy (Fig. 3c-d). This implies that in practice, independently of which *n* strains ultimately coexist in a system out of a pool of *N*, for a given *n*, we should tend to observe such system around the same rank-frequency distribution of strains. Such “optimal” rank-order frequency distribution will display higher variance for low number of coexisting strains (for example *n* = 4), but be more robust for higher *n*, (e.g. *n* = 8), and generally independent of community composition.

While Figure 3 refers to a scenario of low *μ* where co-colonization is dominant, in the Supplementary Figure S5 we explore also the case of a high *μ* tending to the opposite limit of single colonization dominance (μ = 10 is more realistic for a system like pneumococcus serotypes). Strikingly the fundamental feature that diversity increases stability, is preserved. Likewise, the trade-off between probability of reach and stability of coexistence equilibria is also preserved. This means the ecological forces leading to an optimal effective stability in an intermediate entropy range are robust, independently of *μ*. In this limit, we also verify that for the same random structures of interaction coefficients A, typically more strains coexist, an odd number of strains is more likely to coexist (Chawanya and Tokita, 2002), and that their coexistence dynamics are more complex (Box 2).

### The role of *μ* in multi-stability

We revisit the relationship between complexity of the dynamics and ratio of single to co-colonization (*μ* = 1/*k*(*R*_0_ − 1)) in our model, now with simulations over a larger range of *μ*. Using 100.000 random system simulations with *N* = 10 for different values of *μ*, we confirm the pattern observed for *N* = 2 and illustrated for *N* = 4 in Figure 2, that increasing *μ* (via decreasing either *R*_0_ or *k*) increases strain turnover and the complexity of multi-strain coexistence. Computing the number and stability properties of equilibria for each system, we can see that multi-stability of steady-states is very common, especially in the limit of single colonization dominance (*μ* → 0) as shown in Fig. 4a. In contrast, as *μ* increases, multi-stable equilibria become less common but the proportion of systems without any stable steady state increases, pointing to an increase in dynamic complexity for the same *N* and normalized interaction matrix *A* between strains.

**Figure 4:**
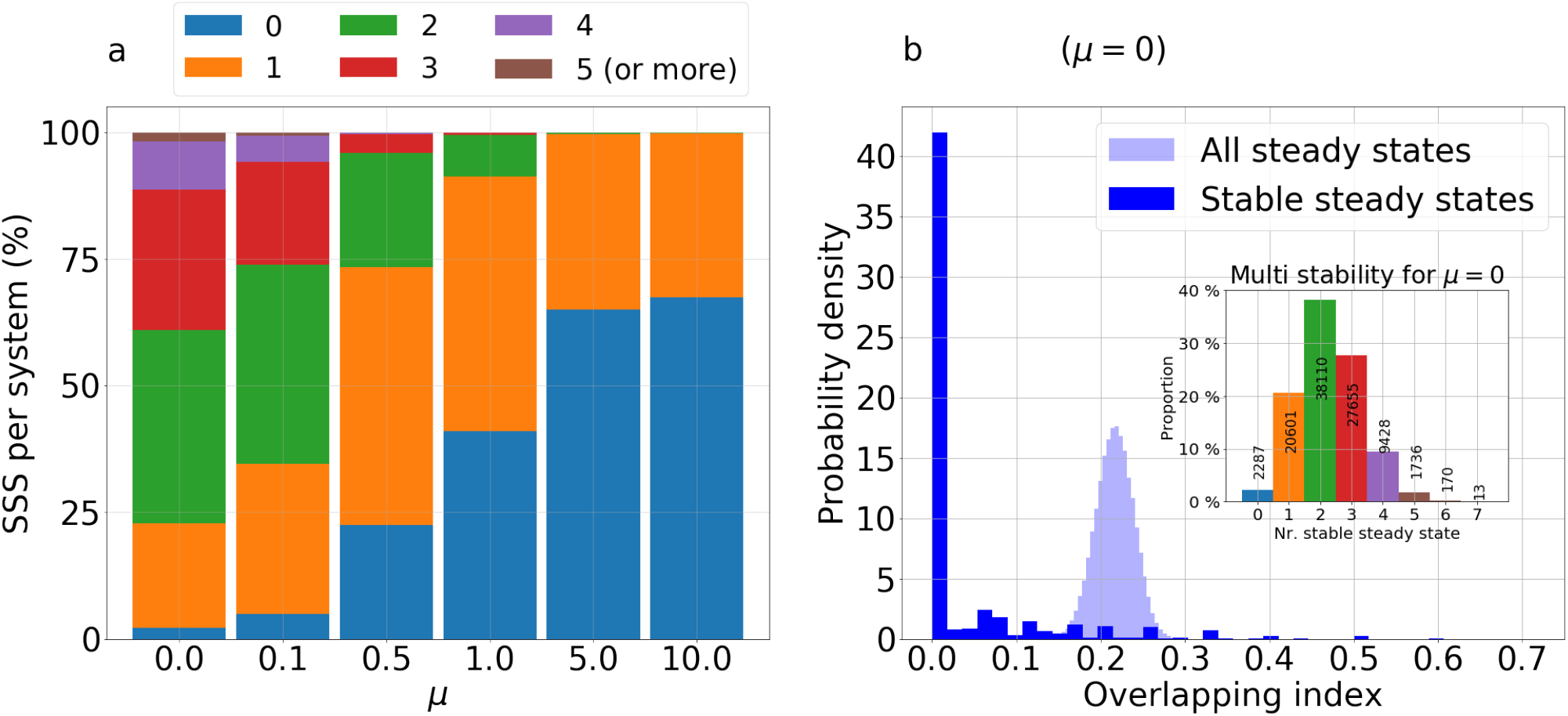
System complexity for large *N* and multi-stability. **a.** For several values of *μ*, 100.000 systems of *N* = 10 strains have been generated (using a normal distribution of *A*). For each value of *μ*, we show the proportion of systems with a given number of stable steady states (in {0,⋯, 6}). As *μ* increases, the proportion of systems without any stable steady state increases, reflecting larger potential for complex coexistence dynamics, where the attractors may be limits cycles, May Leonard trajectories or even strange attractors. For *μ* < 0.5 (high transmission or competition between strains), stable multi-strain coexistence is very probable, and most of the systems have several stable steady states, thus multi-stability is common. **b.** In the particular limit case of *μ* = 0, and for each of the 100.000 systems, we compare the strain overlapping index *SOI* (see Materials and Methods) within *all* steady states and within only the *stable* steady states. We see that without the constraint of stability, two steady states of our system share on average 20% of their coexisting strains, while within the set of stable steady states, this overlapping index is nearly always 0. This shows that in multi-stability, the attractors contain mostly disjoint sets of strains.

In particular, when zooming-in on the multi-stable equilibria of each system, we can quantify how different are the subsets of strains coexisting in each of those equilibria. Defining an index of strain overlap, based on the Jaccard index between two sets (see Materials and Methods), we find that multi-stability is largely characterized by non-overlapping subsets of strains (Fig. 4b), in contrast to the higher average strain overlap (about 20%) between any two generic steady states. This means that depending on initial conditions and founder effects, the same global pool of *N* strains with random co-colonization interactions, can lead to entirely different community composition of strains in different epidemiological settings, an effect that is rather common (60% probability) for values of *μ* around 0.1, but becomes improbable for *μ* = 10.

When quantifying system outcomes for *N* strains, in three broad categories: monostability, multistability, and complexity, with an even higher resolution for *μ*, we find that there are effectively 3 regions (Figure 5): for small *μ*, stable coexistence and multi-stability is more likely, for intermediate *μ* monostability is more likely, and for high *μ*, (i.e. very small *R*_0_ or *k*) complexity, May-Leonard type coexistence and unstable coexistence are more probable outcomes. This result highlights the importance of the environmental gradient in shifting the qualitative dynamics: the more cooperation there is between all strains on average (*k* higher), or the higher the overall transmission intensity (higher *R*_0_), i.e. the lower *μ* is, the weaker the selection imposed by co-colonization asymmetries, and hence stable coexistence (in multiple combinations) is more probable. In contrast, when *μ* increases, either via more competition between strains on average (*k* lower), or lower transmission intensity (*R*_0_ closer to 1), the stronger the importance of asymmetries, and thus the stronger the selection imposed by co-colonization trait differences between strains. In that extreme, because the strains are so inter-dependent, coupled more strongly via the available dynamic host resources in the single infection compartment (implying dynamic niche creation and destruction along at most *N* axes), the probability of unstable coexistence between them is largest. In one extreme, we expect low variability in a setting over time, in the other extreme we expect high variability and complex strain turnover over time.

**Figure 5:**
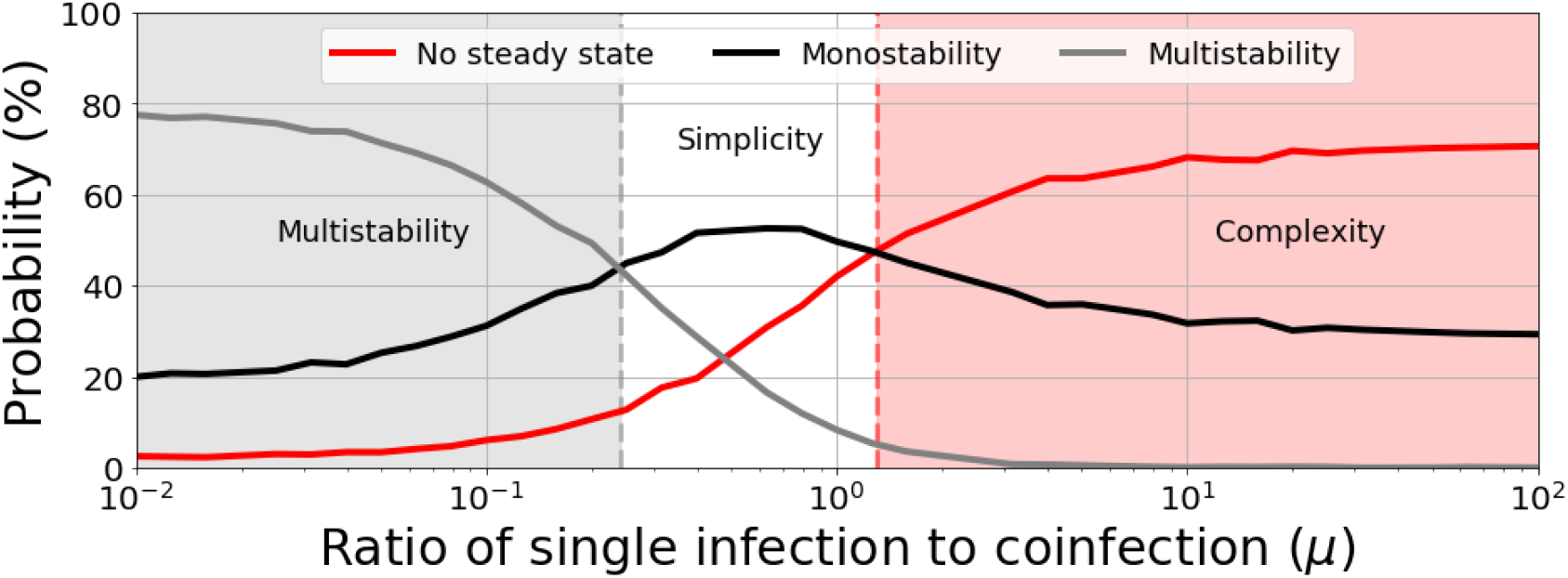
Dynamic regimes as a function of singe to co-colonization ratio *μ*. We plot the number of steady states as a function of *μ* G [0.01,100] for many randomly-generated interaction matrices A. Since *μ* depends on *R*_0_ and *k*, we kept their combination implicit, and randomly sampled the rescaled interactions matrix *A* = (*α_ij_*). For any system (*A*), if there are two or more stable steady states, the system is considered multistable and the dynamics depend on the initial conditions. If there is only one stable steady state the dynamics can be considered simpler, in principle every system with a unique global attractor is monostable, but there may be other more complex unique attractors like stable limit cycles. If there is no steady state, the dynamics are more complex, meaning the attractors are necessarily neither stationary like limit cycles, heteroclinic cycles or even strange attractors. For the simulations, we used a total strain pool size of *N* = 10, and for each value of *μ* we generated randomly 1000 matrices A leading to 1000 different interaction systems for each value of *μ*. For a given value of *μ*, we then counted the proportion of systems with no stable steady state (red line), exactly one stable steady state (black line) or several stable steady states (grey line). For a small *μ*, very few systems have zero stable steady states, while nearly 80% of them are multistable. For a large value of *μ* the multistability is nearly impossible while more than 70% of interaction systems having no steady state and thus displaying a complex dynamics with no stationary attractors.

Constructing multi-strain communities with random co-colonization interaction strengths (in normalized matrix *A* form), we find that the global epidemiological quantity 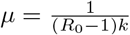, namely the single to co-colonization ratio modulates regimes of system behavior and provides insights on the diversity-stability debate (McCann, 2000). For a fixed *μ* and a given set of interaction coefficients between strains however, as their frequencies change epidemiologically, the epidemiological dynamics and the evolutionary dynamics interact. Evolution happens on two levels in this system. First, frequency dynamics (*z_i_*) can be linked with selective dynamics on mean interaction in co-colonization trait space (see (Madec and Gjini, 2019)) where *k_effective_* = *k* + *εq(t)*. Secondly, another trait changing among community members is mean invasibility of the system, *Q*, in mutual invasion fitness space (see Box 1). These two quantities reflect the changing mean fitness landscape, and depend on frequency dynamics as quadratic terms involving the product between strain pair frequencies over time. When varying *μ*, as the multi-strain selective dynamics unfold in different ways, so do the evolutionary dynamics of mean traits in the system (Box 3).

#### Box 3. Stability-diversity-complexity and evolutionary dynamics of interaction traits

In multi-strain communities with random co-colonization interaction strengths (*A*), the global epidemiological quantity 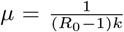, describing the single to co-colonization ratio modulates regimes of system behavior, shaping its stability, complexity and diversity properties.

Together with epidemiological dynamics, evolution happens in real time on two traits in this system: i) mean interaction in co-colonization trait space (*q*) where *k_effective_* = *k* + *εq*(*t*), ii) mean invasibility of the system *Q* in mutual invasion fitness space (Eq. 3).

Evolutionary dynamics in the microbial community reflect the changing mean fitness landscape, and depend on frequency dynamics (Madec and Gjini, 2019) as follows:

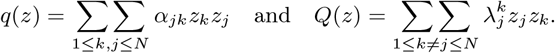

Notice that *q* increasing means the system tends to increase cooperation on average over time, whereas *q* decreasing means the multi-strain community tends to increase competition over time. Similarly *Q* increasing implies the system becomes less invadable over time, and *Q* decreasing means the community of strains becomes easier to invade from outsider strains. As frequency dynamics can be complex, so can mean trait dynamics.

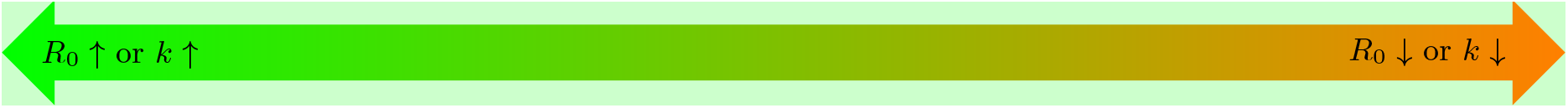

- **Cooperation or high transmission** (*μ* **low**)

– Few species coexist
– Multistability is common
– Simple coexistence (fixed point)
– Low variability over time in one population
- Mean trait evolution on 2 levels: *q* and *Q* over time

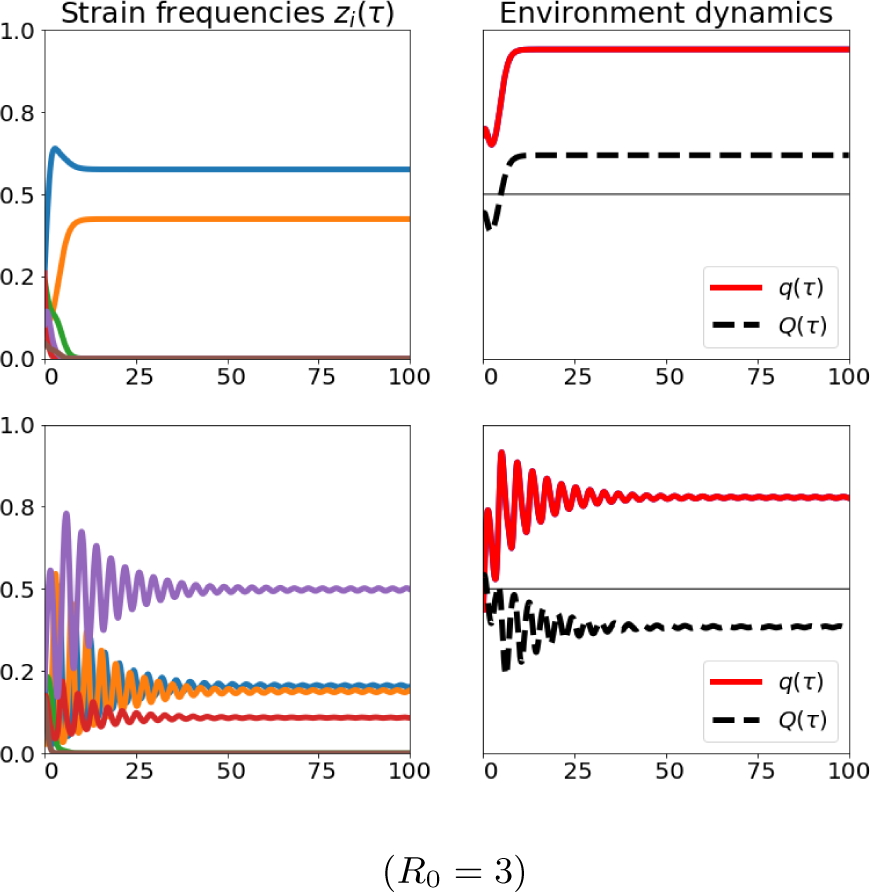
- Either competition or cooperation can be selected
- Evolution of “specialist” communities
- High diversity between host populations
- Competition or low transmission (*μ* high)

– Many species coexist
– Unstable coexistence common (no multi-stability)
– Complex attractors (limit or heteroclinic cycles)
– High variability over time in one population
- Mean trait evolution on 2 levels: *q* and *Q* over time

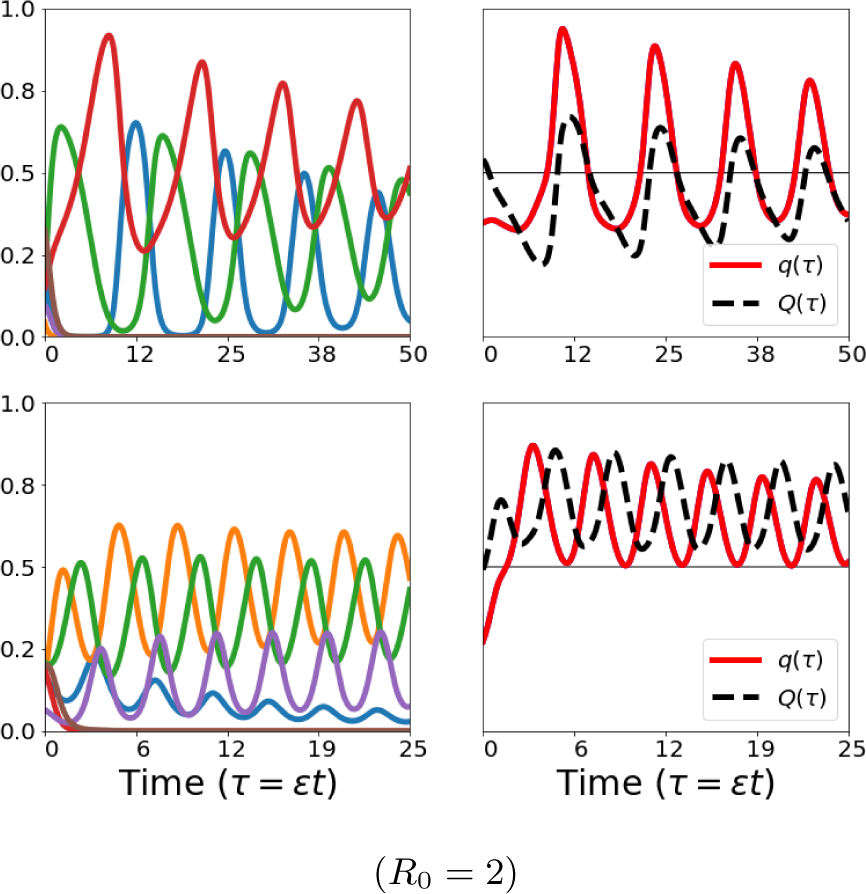
- Competition and cooperation fluctuate!
- Evolution of “generalist” communities
- Lower diversity between host populations

In particular, in the *μ* → 0 limit, where multi-stability is common, systems can evolve towards more competition or more cooperation in co-colonization depending on initial conditions. Conversely in the other limit *μ* → ∞, where complex oscillatory coexistence is more probable, systems preserve polymorphism in interaction trait space, with alternating periods between competitors and cooperators. This result highlights two routes to coexistence or maintenance of biodiversity: little variability within one population but diversity between populations in the first case, and more variability within one population but low diversity between populations in the second. Notice also that increase in mean resistance to invasion of the multi-strain system (*Q* evolutionary dynamics) does not necessarily always correlate with increased cooperation or competition (higher or lower *q*), and in the oscillatory regime the two can also be synchronous and asynchronous (Figure S6).

### The *R*_0_ vs. *k* trade-off in *μ* and the stress gradient hypothesis

The structure of a multi-strain system is the result of complex interactions among different members, where the relative importance of each interaction can also be a function of the global environmental context, and of the biological interactions among the rest of the community. This context-dependence in effects of ecosystem engineers for the rest of the community has long been highlighted in ecological systems (Hastings et al., 2007). In our model, we have seen that strains can mutually engage in more or less competitive interactions and more or less cooperative interactions, and differential abilities in such traits allow each strain to potentially act as an engineer. Thanks to this similarity assumption for dynamics decomposition, we can disentangle very clearly the effect of mean interaction *k* and the variation around that mean in what concerns intra-specific and inter-specific interactions, *A* = (*α_ij_*), and we can also see how mean ‘environmental’ variables contribute to relative fitness between system members λ_*ij*_. Until now, we have seen that in our system, the ratio of single to co-colonization *μ* = 1/(*R*_0_ − 1)*l*) is a key modulator of the qualitative complexity of the dynamics (Figure 5, Box 2). Thanks to the simplicity of this formula, it is clear that both transmission intensity *R*_0_ and mean interaction coefficient in cocolonization *k* decrease this ratio, and the only way for *μ* to be held constant, is if Ro and *k* are traded-off against one another. In other words, in order to keep *μ* constant when transmission opportunities for all strains are reduced, the mean interaction coefficient in the system must increase, i.e. if all strains cooperate more in co-colonization. And viceversa, if the environment gets more favourable, i.e. *R*_0_ increases, then *μ* can be kept constant only if *k* decreases, thus if all strains increase competition. Such exact quantitative finding in our model makes an explicit link with the stress-gradient hypothesis (SGH) postulated in ecology (Bertness and Callaway, 1994; Callaway and Walker, 1997). The SGH predicts that positive interactions should be more prevalent in stressful environments, while more favourable environments should favour competition. This hypothesis has been supported and tested mainly in plant systems (Callaway et al., 2002; Eränen and Kozlov, 2008; Pugnaire and Luque, 2001), and more recently in microbial communities (Fetzer et al., 2015; Hoek et al., 2016; Lawrence and Barraclough, 2016; McCluney et al., 2012; Piccardi et al., 2019).

In our system, which is an epidemiological system at basis, but with microbial interactions embedded in co-colonization dynamics, we can verify via the single-to co-colonization ratio, an interpretation of the stress-gradient hypothesis. In order for a set of species with a fixed set of normalized interactions (A=(*α_ij_*)) between them, to maintain a given configuration of coexistence (*μ*) under changing environmental conditions (e.g. *R*_0_), the only way to do it is by universal adaptation in mean-field interaction, increasing competition in favourable environments and increasing cooperation in harsher environments, respectively (see Figure 6). However, it is worth noting that even if *μ* is kept constant, i.e. even if the species respond to harsh environmental changes by increasing cooperation, and thus preserve their qualitative equilibrium configuration, the speed of their collective dynamics may still be affected, typically reduced with *R*_0_, although the absolute magnitude will depend on whether and how much they are mediated through altered transmission rate vs. altered duration of carriage (see Supplementary Figures S3–S4). In summary, while the fitness landscape is shaped by the phenotypic distribution of all the community members, as the external environment changes, the relative balance of their interactions may shift, and one way to keep that balance in check would be by quick adaptation of the mean.

**Figure 6:**
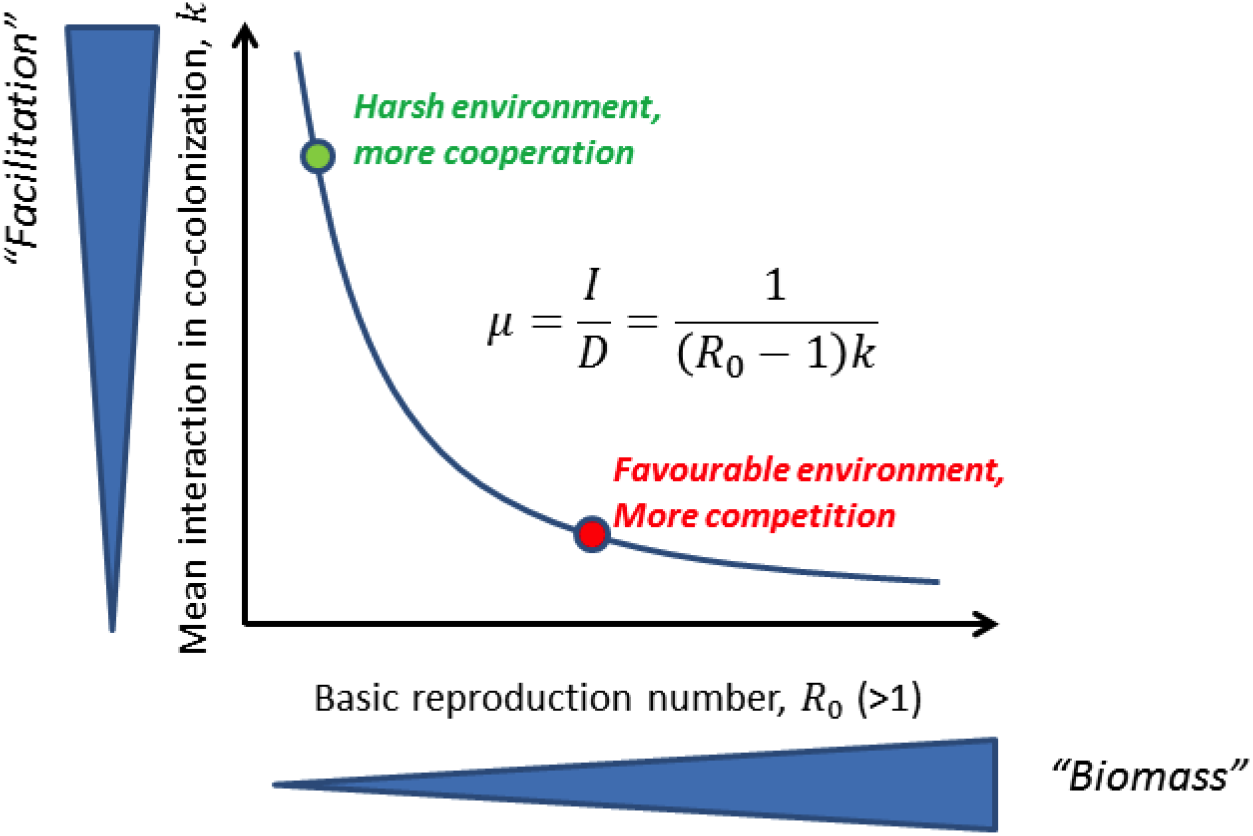
Transmission intensity *R*_0_ and mean interaction in cocolonization determine *μ* in a tradeoff manner. While the type of equilibrium where the system will tend to, is strongly driven by *μ* (the ratio of single to co-colonization), hence directly by *R*_0_ and *k*, it is clear from the expression 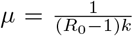 that the only way for *μ* to be held constant, is if *R*_0_ and *k* are traded-off against one another. This principle reflects the expectation from the stress-gradient hypothesis (SGH): when transmission opportunities for all strains are reduced (harsher environment), the mean interaction coefficient in the system must increase, i.e. if all strains should cooperate more in co-colonization, to keep the same *μ* and hence qualitative coexistence regime. And viceversa, if the environment gets more favourable, i.e. *R*_0_ increases, then *μ* can be kept constant only if *k* decreases, thus if all strains increase competition. Note that keeping *μ* fixed only preserves the qualitative nature of the dynamics between strains, but does not guarantee that its speed Θ will remain constant. Changes in global *R*_0_ and *k* and the underlying mechanisms giving rise to such changes, will be important drivers of changes in selection speed.

## Discussion

A central question in microbial ecology is whether community members compete or cooperate with one another, and how such interactions mediate community stability, resilience and function. In our model, we study an epidemiological multi-strain system, where members interact with each other via altered susceptibilities to cocolonization, which broadly include both competition and facilitation. By clearly delineating the role of mean interaction coefficient, basic reproduction number *R*_0_, and biases in pairwise coefficients relative to the mean, we obtain a model reduction for strain frequency evolution for any *N*. The questions we addressed within such a system are similar to a long-standing ecological quest on multispecies dynamics (May, 1972; Pascual et al., 2006; Serván et al., 2018): what governs coexistence regimes, diversity and stability in such systems?

Although we do not specify the molecular mechanisms that can mediate positive or negative interactions between strains in co-colonization (Dawid et al., 2007; Leggett et al., 2014; Lysenko et al., 2010; Riley and Gordon, 1999; Shen et al., 2019) quantitatively we can predict very important features of system behavior with a simple mathematical framework derived under quasi-neutrality and strain similarity assumptions (Madec and Gjini, 2019). This model captures selective dynamics between strains over long time, and coincides with an instance of the replicator equation (Hofbauer and Sigmund, 2003), parametrized in our case, in terms of pairwise invasion fitnesses between strains, a central quantity in adaptive dynamics (Geritz et al., 1998). Studying this equation and its epidemiological consequences in detail, here we find that there is a critical ratio that tunes asymmetry and dynamic regimes in such multi-strain interacting system: namely the ratio of single to co-colonization. This ratio *μ*, given by the inverse of the product of basic reproduction number and mean permissiveness to co-colonization *μ* = 1/((*R*_0_ − 1)*k*), increases or decreases the importance of asymmetry in interaction between strains as they encounter each-other epidemiologically in resident-co-colonizer combinations, and lends support to the context-dependence (Coyte and Rakoff-Nahoum, 2019) of relative fitnesses in a coupled microbial community.

In our epidemiological system, individual strain frequency evolution over long times depends on how that strain interacts with other strains, but also on the total net interaction between extant strains. Such global, symmetric and temporally-varying, environmental feedback on all strains emerges as a key feature of this cocolonization model, and can be used to summarize the evolution of diversity in the system. It will tend to increase when more strains are stably maintained over time, and it will tend to zero in the extreme cases of competitive exclusion and chaos. Each strain is competing for susceptible hosts, and for (at most) *N* singly-colonized hosts that can be co-colonized. These *N* types of available hosts are the resources upon which the strains can grow epidemiologically, but these resources are not independent from each other. With each additional strain, the landscape of interactions can change dramatically for any system composition, because each additional strain offers a new type of resource upon which the competition of the residents can be manifested, while also modulating directly the relative success of those strains by taking over space from their colonization. This dynamism embedded in the inter-connected and inter-dependent system is the source of dynamic complexity, recognized generally in biological games (Nowak and Sigmund, 2004), and here uncovered and quantified in an explicit epidemiological context.

We find that for small values of the ratio between single and co-colonization *μ*, which means high transmission intensity or high cooperation among strains on average, the system dynamics is characterized by multi-stability, where typically non-overlapping strain subsets can coexist, depending on initial conditions. This result is consistent with earlier multi-strain SIRS models with cross-immunity, that in order to minimize competition and maximize chances of persistence, polymorphic pathogens self-organize in non-overlapping subsets (Gupta and Anderson, 1999). Here, within a much simpler, SIS framework, we find a similar principle applies, despite the interactions between strains being mediated via altered susceptibilities to co-colonization, without persistent immune memory.

Our study brings together several themes of interest across many multi-type systems shaped by interactions and higher-order feedbacks between their members. Many fields including ecology, geophysics, hydrology and economics are calling attention on critical transitions, which occur when natural systems drastically shift from one state to another (Scheffer et al., 2012). Critical transitions in the epidemiology of infectious diseases are of relevance to the emergence of new pathogens and escape from control, such as vaccines. The critical transitions analyzed in this paper relate global and mean-field environmental variables to the manifestation of competitive hierarchies between multiple strains interacting in co-colonization. We have made explicit how a gradient emerges from the epidemiological ratio of single to co-colonization, and tunes effectively the diversity, stability and complexity of the coexistence between strains. Such gradient can mediate critical transitions in collective dynamics, when the normalized interaction coefficients between members are held fixed. These transitions may underlie and potentially enhance (or counteract) efforts to control and eliminate multi-type infectious pathogens, as via vaccines or drugs, or in the face of climate change.

In a recent theoretical study focused on malaria, without explicitly modeling multiple strains and host immune memory, superinfection (concurrent multiple infection) has been shown to consistently create tipping points that give rise to hysteresis in responses to control or seasonal variation in vector abundance (Alonso et al., 2019). Our present work, developed in the framework of many interacting strains, lends support to a similar perspective, but more generally relevant to high-dimensional conservative SIS systems and coexistence regimes, rather than prevalence tipping points. Cooperation and competition among strains, studied here under the same integrated context, appear as two sides along a continuum for the system, which particular strain compositions or environmental drivers (seasonality, general host immunity, population turnover) may tip towards one or the other extreme. We find that when *μ* tends to favor co-colonization, for example in the limit of more facilitation between strains on average, the system tends to multi-stability and stable coexistence of a few strains in simple dynamics. In contrast, when *μ* tends to favor single colonization, for example in the limit of more competition between strains on average, the system tends to more complexity and unstable equilibria (Fig. 5), but coexistence of more strains becomes possible (Fig. S1). Although at first sight this may seem to suggest, somewhat contrary to previous expectation from lower-dimensional models (Chen et al., 2017; Hébert-Dufresne and Althouse, 2015), that average cooperation in co-colonization is stabilizing and average competition is destabilizing, our result should be read in the broader view of the underlying resource dynamics shaped dynamically by *N* strains, and the opportunities for small trait differences between them to get manifested in a homogeneously mixing host population. When contact structure heterogeneities are accounted for, it would be interesting to test, how much of this behavior will still be preserved.

Taken together, our results show that even though differences in pairwise interactions can be small and random, as expected for closely related strains, a large community of such strains can collectively coexist, and self-organize, albeit often in complex and unstable fashion, simply thanks to the critical interplay between global transmission potential and mean permissiveness to co-colonization. We found that the requirement of being simultaneously stable and feasible tends to push coexistence regimes toward intermediate diversity, independently of precise composition. We argue that the stress gradient hypothesis (SGH) (Bertness and Callaway, 1994; Callaway and Walker, 1997) can be used as a framework to help interpret the critical role of the single-to colonization ratio in shaping epidemiological multi-strain systems.

Finally, this model makes several predictions which can be tested empirically. First, the invariant principles in the slow time scale (Box 1) suggest that dominance patterns in single and co-colonization of particular strains should be the same. The other finding that stable (multi-stable) coexistence between types through cocolonization is more likely when mean interactions tend towards cooperation, and that a single stable coexistence between types becomes more probable at intermediate values of *μ*, and ultimately only unstable coexistence is possible for large values of *μ* could be tested in endemic multi-strain pathogen systems. For example, empirical data in polymorphic *S. pneumoniae* bacteria, have been consistent with estimates of about 90% mutual inhibition between co-colonizing serotypes (Gjini et al., 2016; Lipsitch et al., 2012), and *R*_0_ values around 2. This implies that for this system, *μ* ≈ 10, and multi-stability is highly unlikely but a single stable equilibrium point is almost as likely as complex unstable coexistence, which may reconcile the secular trends observed in some settings consistently over many years (Ekdahl et al., 1998; Feikin and Klugman, 2002; Fenoll et al., 1998). Such secular trends can interfere with vaccine introductions and need to be accounted for when estimating impact (Moore, 2009; Moore and Whitney, 2008). Our model makes explicit predictions about the timescale and qualitative aspects of such secular trends (Box 2, Fig. S4), under the plausible assumption that they are driven by co-colonization interactions.

The consistency of the optimal diversity (entropy) configuration for a given number of strains, is a model prediction that has in fact been observed empirically for pneumococcus serotypes across geographical settings (Hanage et al., 2010). While the serotype composition has changed in different populations before and after vaccination, the rank-frequency distribution of serotypes, related to entropy in our model, has been conserved. Our model, offers a new perspective on such empirical observations, in terms of a null expectation for how such pattern emerges and is maintained, without needing to invoke strain-specific transmission rate dominance or duration of carriage dominance (Weinberger et al., 2009), but simply based on co-colonization interactions between serotypes, which when random balance out in a particular optimal entropy configuration. Since this optimal rank-order abundance distribution depends on the context *μ*, model expectations for such dependence can be tested with data from different *R*_0_ settings. Finally, the link with the stress gradient hypothesis, suggests that to preserve a certain “optimal” single-to cocolonization ratio (optimal complexity/coexistence balance), independently of community size *N*, facilitation in co-colonization between microbial strains should be more common in epidemiological settings with low prevalence, and this is an interesting prediction to test in the future. It becomes intriguing to verify, beyond pneumococcus, to what extent this model and its insights (Box 3) can be used as an analytic backbone to revisit and interpret multi-strain dynamics in other systems of relevance for public health, for example influenza (Yang et al., 2019), dengue (Mier-y Teran-Romero et al., 2013), malaria (Alonso et al., 2019; Gupta and Maiden, 2001) or human papilloma viruses (Murall et al., 2014).

Disentangling multi-strain interactions and their role in community function at the epidemiological level remains challenging, but can be made more accessible analytically using frameworks such as the one proposed here. With the simplicity and deep insights afforded by this model, we can address better the role of mean fitness of the microbial system as a whole, trait variance, and the role of environmental gradients for stabilizing vs. equalizing forces in biodiversity.

## Methods

### N-strain SIS model with co-colonization

The model is given by the following equations:

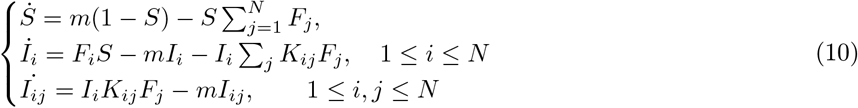

where 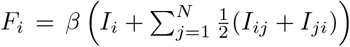 gives the force of infection of strain *i*, and *S*, *I_i_* and *I_ij_* refer to the proportion of susceptible hosts, singly-colonized by strain *i* and co-colonized by strains *i* and *j*. Notice that *S* = 1 − Σ*I_i_* + *I_ij_*, thus the dimension of the system is effectively *N* + *N*(*N* − 1)/2. A key assumption in the model is that hosts co-colonized with different strains *i* and *j* transmit either with equal probability. In the above notation *m* = *γ* + *r*, is a population turnover rate, encapsulating both clearance rate *γ* of colonization episodes and recruitment rate of susceptible hosts *r* (balanced by natural mortality rate *r* = *d*).

This model has been described in detail elsewhere (Gjini and Madec, 2017) for the case of *N* = 2. The model allows for each strain to interact differently with other strains upon co-colonization, altering the suceptibility of an already-colonized host to incoming strains. The magnitude and type of such interactions is described by the matrix *K*, where values *K_ij_* above 1 indicate facilitation between strains, and values of *K_ij_* below 1 indicate inhibition or competition between strains. We do not make any specific assumptions on the mechanisms underlying such interactions, but a key feature of this formulation is the explicit incorporation of intra-strain and inter-strain interactions. The derivation of the slow-fast dynamics decomposition lies on the assumption that the variance of such interaction coefficients is low, thus deviation from neutrality of *K_ij_* with respect to a reference *k* is small. In our particular simulations here, we assume a normal distribution for such interaction coefficients. We can write every *K_ij_* as: *K_ij_* = *k* + *εα_ij_* under the assumption that *ε* is small.

Although many different parametrizations are possible, we define 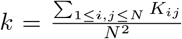 as the statistical mean of the co-colonization interaction coefficients between all possible strains in the available pool. We define the deviation from neutrality *ε* as the standard deviation of (*K_ij_*)_1≤*i,j*≤*N*_. This leads to the normalized interaction matrix

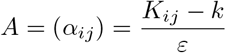

to have the same distribution as *K*, but with mean 0 and variance 1. We use analysis and simulations to understand the behavior of the epidemiological model for different assumptions on the co-colonization interaction matrix, and for different system size *N*. We use the reduced system (of *N* equations) for dynamics on the slow timescale *εt* (Madec and Gjini, 2019) to obtain the steady-states of the system and analyze their properties and stability.

### Strain overlap index (SOI) for two coexistence equilibria of the same system

Consider a set *S* = {*E*_1_, *E*_2_,⋯, *E_p_*} with all steady states of a given system (interaction matrix), where *E_j_* ⊂ {1,⋯,*N*} and 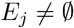. In our applications, *E_j_* is the index of feasible strains (with *z_i_* > 0) in the steady state number *j* of a given system of *N* = 10 strains. We note 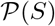 the set of each subsets of *S*. As instance, if *S* = {*E*_1_, *E*_2_} then 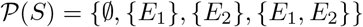. Recall that 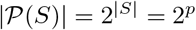.

We denote the strain overlapping index by SOI and define it as a mean over the Jaccard indices calculated for each pair of coexistence equilbria:

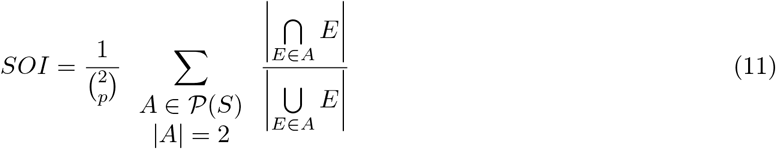

## Acknowledgement

EG acknowledges support by the University of Tours and EURAXESS for an invited researcher stay at IDP, in Tours during 2019, which was instrumental to complete this collaborative work.

## Supporting information

### S1 Supplementary figures

**Figure S1:**
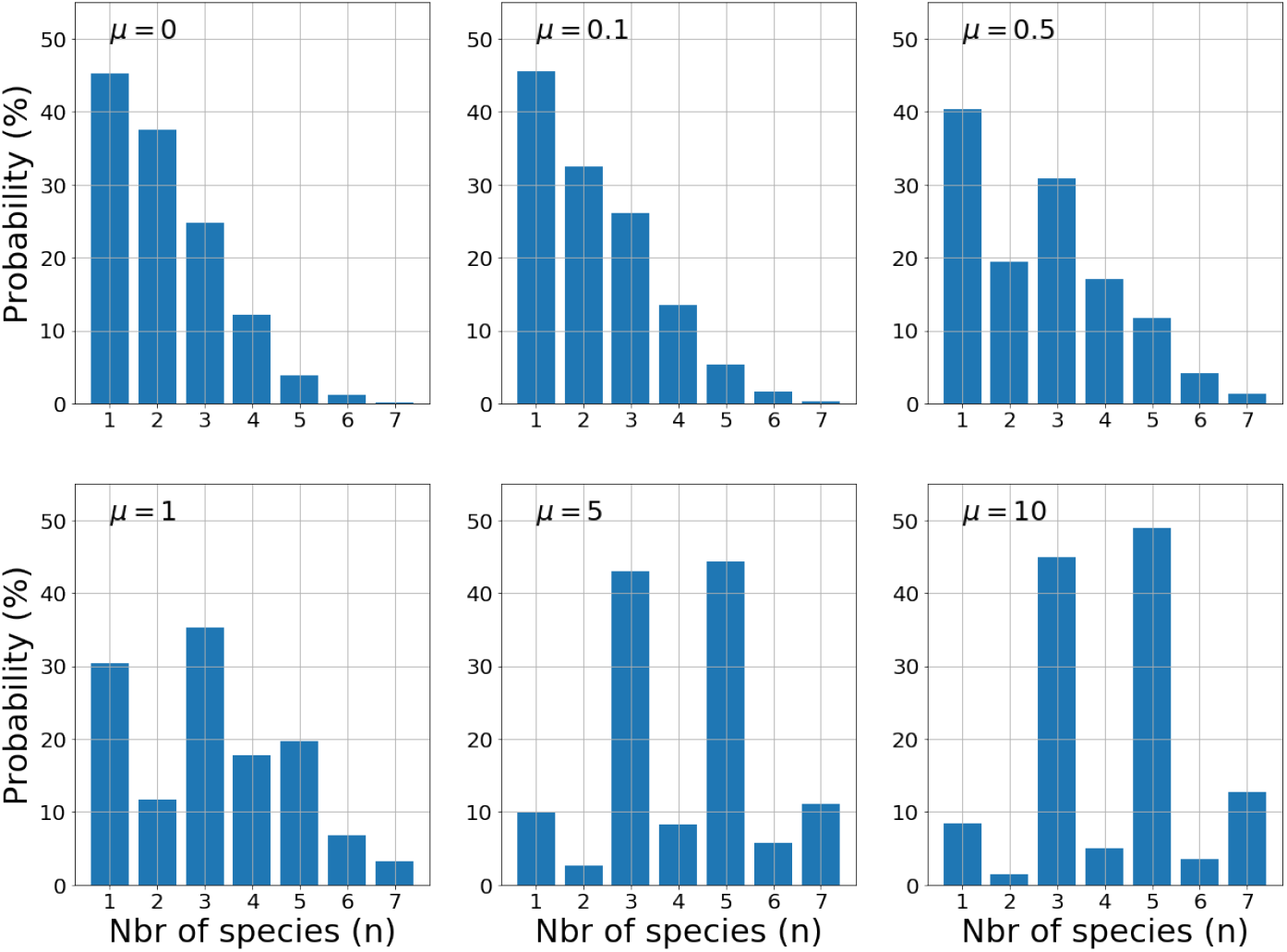
Probability for stable *n*-strain coexistence out of a pool of *N* = 10 strains. We plot the distribution of the number of coexisting strains *n* for 6 values of *μ*. For each value of *μ*, we generated 100 000 matrices *A* (normalized interactions between strains) and computed their stable steady states. The data are the same as those in the figures 3 and 5. We find that the distribution of *n* is very dependent on *μ*. In general, the probability is hard to compute explicitly but the limit *μ* → ∞ is explicitly known using the fact that the pairwise invasion fitness matrix becomes skew symmetric in that limit (see Box 2). One remarkable fact is that for small *μ* the probability of finding a stable steady state is greater for *n* = 1 and decreases very quickly with number of strains *n*, while for large values of *n*, the distribution looks like a binomial distribution centered at 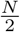 for odd values of *n*, and zero for even values of *n*. Hence, the probability to have a large number of strains coexisting at a given steady state is small for small *μ* and increases for large values of *μ*.

**Figure S2:**
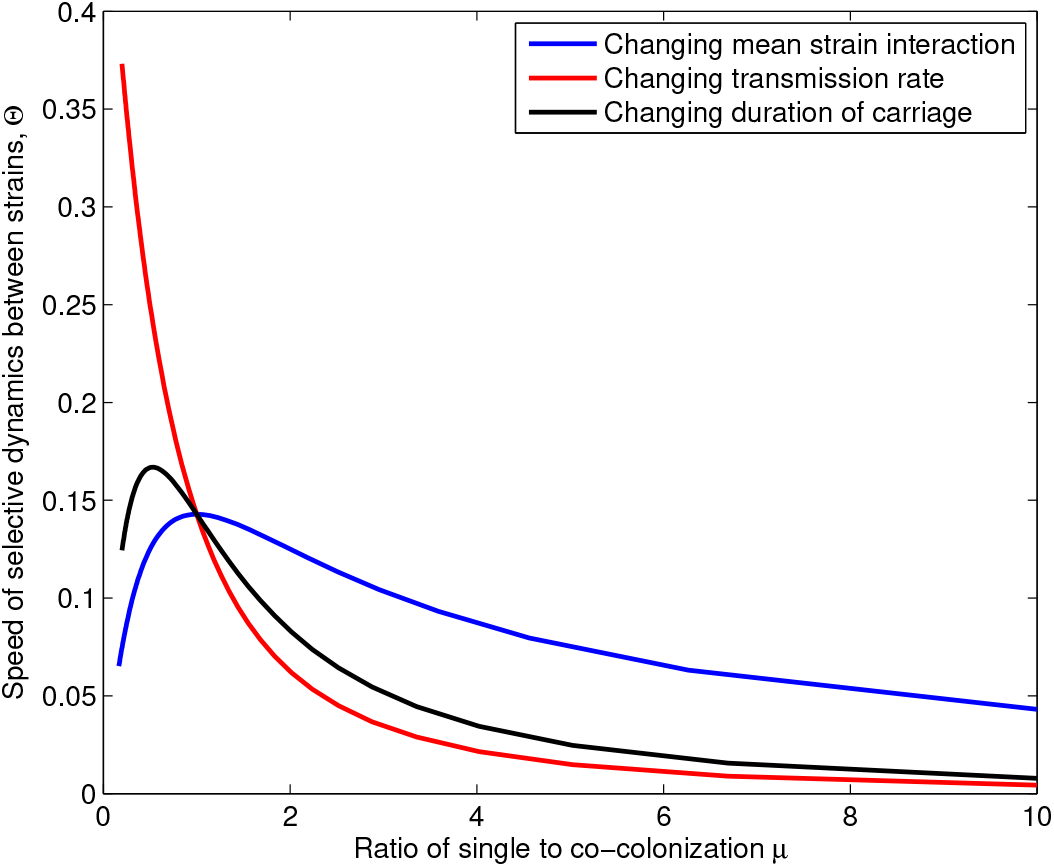
Speed of dynamics depends on *μ* and other parameters. The ratio of single to co-colonization *μ* can vary in three different ways. When *k* is varied: *β* = 2, *m* = 1, *R*_0_ = 2, *μ* decreases with *k* (blue line). When *R*_0_ is varied: via *β*: *k* = 1, *m* = 1, *μ* decreases with *R*_0_ (red line). When *R*_0_ is varied via *m*: *k* = 1, *β* = 1, *μ* again decreases with *R*_0_ (black line). The effects of the same change in *μ* on the parameter 0 are different for these three different cases.

**Figure S3:**
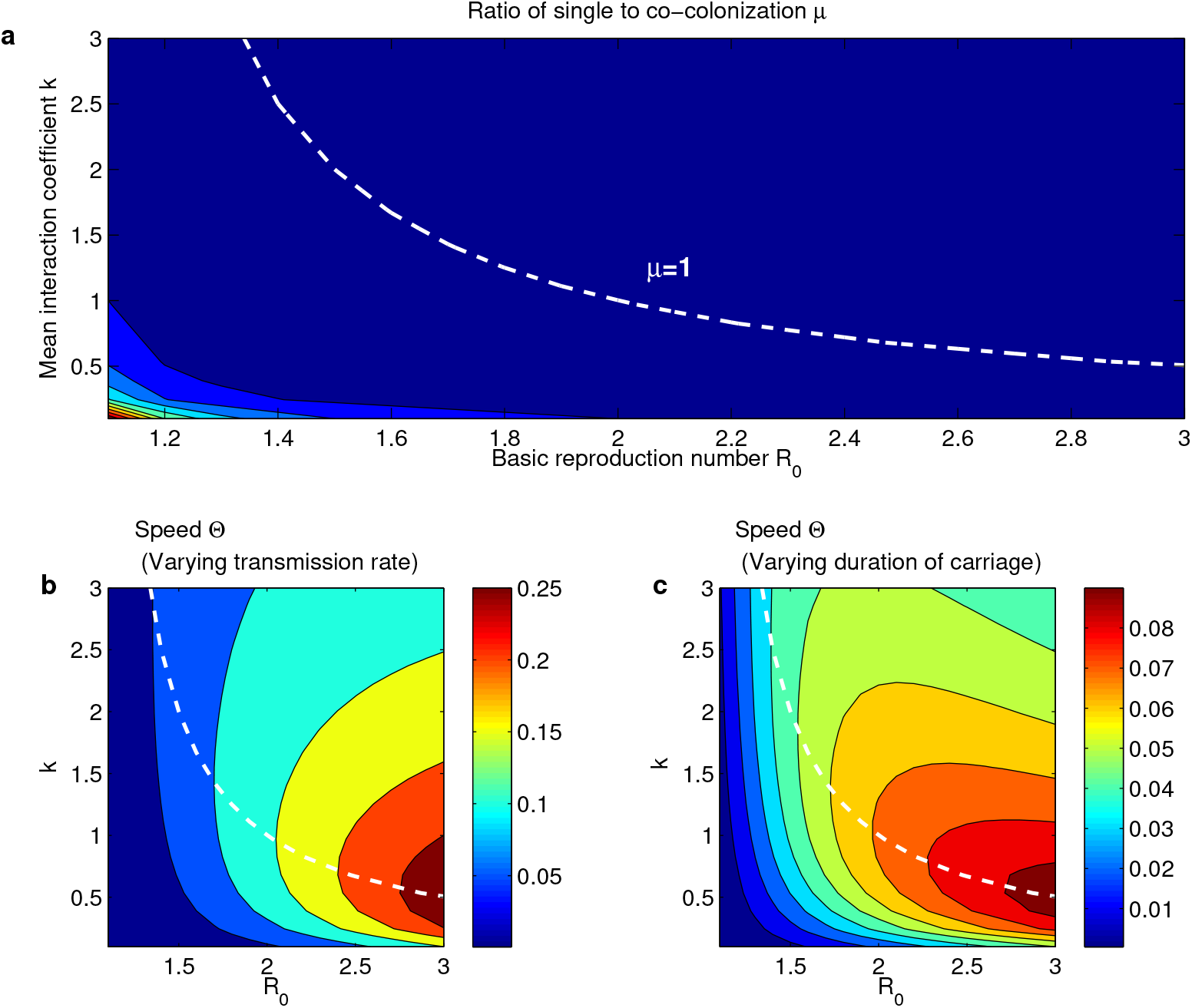
Keeping *μ* constant via the trade-off *R*_0_ – *k* can have different quantitative effects on the speed of the dynamics depending on epidemiological drivers. a. Relationship between *R*_0_ and *k* in determining *μ*. b) Assuming fixed *m* =1, *β* = *R*_0_*m* is varied to give rise to variation in a). c) Assuming fixed *β* =1, *m* = *R*_0_/*β* is varied to give rise to variation in a). The same change in *μ* can lead to different effects on Θ in these two cases.

**Figure S4:**
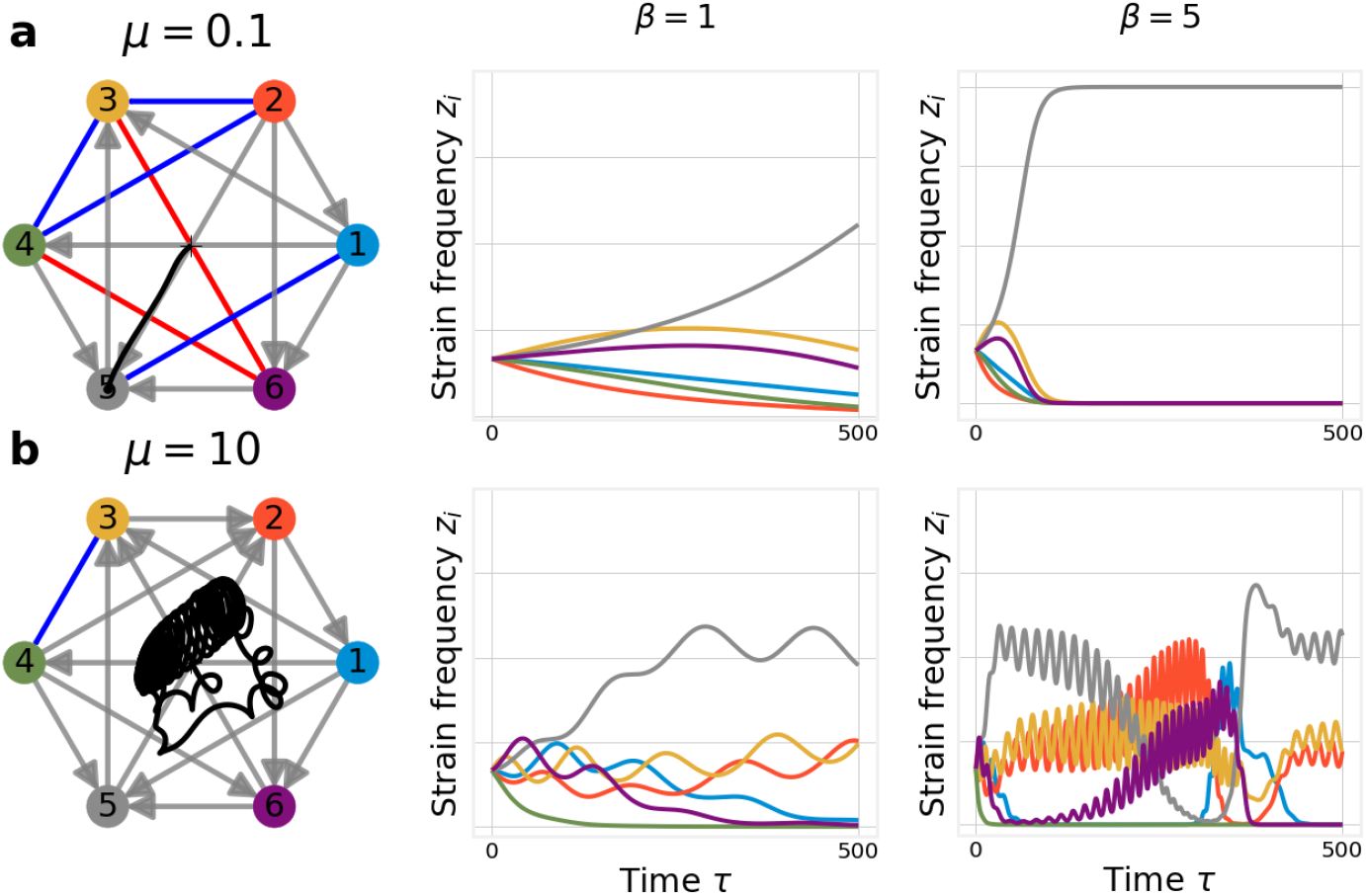
Keeping *μ* constant preserves the same qualitative dynamics in principle but there are timescale effects. For two values of *μ* (rows a and b) we compute the dynamics and we show the multi-strain frequency trajectories for two different choice of transmission rate *β* (columns 2 and 3). *μ* and *β* being fixed, we have *k* = 1/*μ*(*R*_0_ − 1). For the normalized matrix of interactions *A* we assume the same matrix in each figure

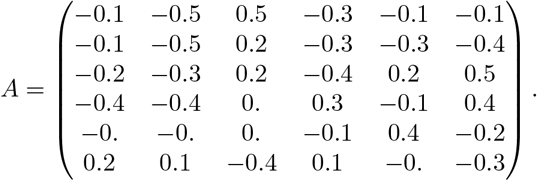 We assume *m* = 0.5 (clearance + natural mortality rate) so that *R*_0_ = 2*β* and a maximum time for simulation *T_max_* = 500. For each row the qualitative dynamics remain the same as shown on the graph. However, the speed of this dynamics depends on *β*. **a.** On the first row *μ* = 0.1, we have a competitive exclusion scenario. Only strain 5 persists. When *β* = 1 (and then *k* = 10 to compensate via the trade-off for *μ* constant) all the strains are still present in the system at *T_max_*. For *β* = 5 (then *k* = 100/9), dynamics are much quicker and selection of finally only one strain occurs very rapidly. The values of Θ are respectively: 0.043, and 0.39. **b.** In the second row *μ* = 10. For this value of *μ*, the same matrix *A* yields more complex dynamics in general, in line with the result that qualitatively the dynamics are determined by *μ*. However, when *β* = 2, the dynamics seems to approach a cycle with dominance of strain 5 (grey) and extinction of strains 1, (blue) 4 (green) and 6 (purple). For *β* = 10 (so *k* = 1/90), the dynamics over the same timescale show a very complex behavior. First strain 6 appears to come back, and 5 goes to extinction, then strain 1 comes back again and ultimately strain 5 dominates in coexistence with 2 and 3. The values of Θ respectively are: 0.043, and 0.39. Thus increasing transmission rate has increased the speed about ten fold.

**Figure S5:**
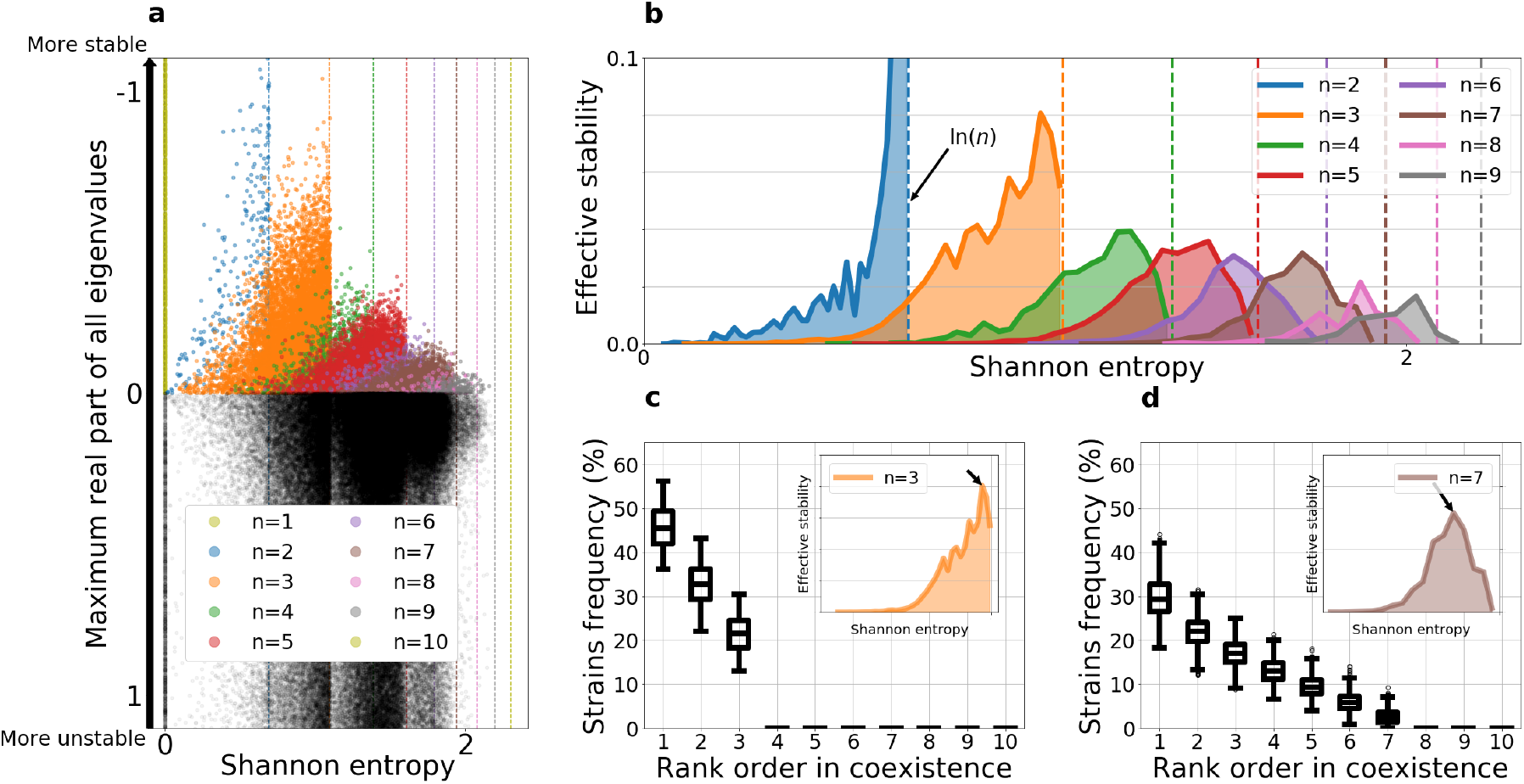
Diversity-Stability relationship for *μ* = 10. This figure is similar to figure 3 but for *μ* = 10 (which is more realistic for pneumococcus). **a.** Relation between Shannon entropy (x-axis) and stability (y-axis).for difference number of *n* coexisting strains within a pool of *N* = 10. Due to this large value of *μ* = 10, it is very rare to observe an even number of coexisting strains *n* (see figure S1). **b.** However, the shape of the effective stability continues to be similar to the case *μ* = 0.05 for both even and odd *n*. For *n* ≥ 3, there is an “optimal diversity” which leads to the optimal balance between feasibility and stability of the steady state. **c. and d.** To be consistent, we illustrate the rank order at the “optimal diversity” for the odd number of coexisting species *n* = 3 and *n* = 7.

**Figure S6:**
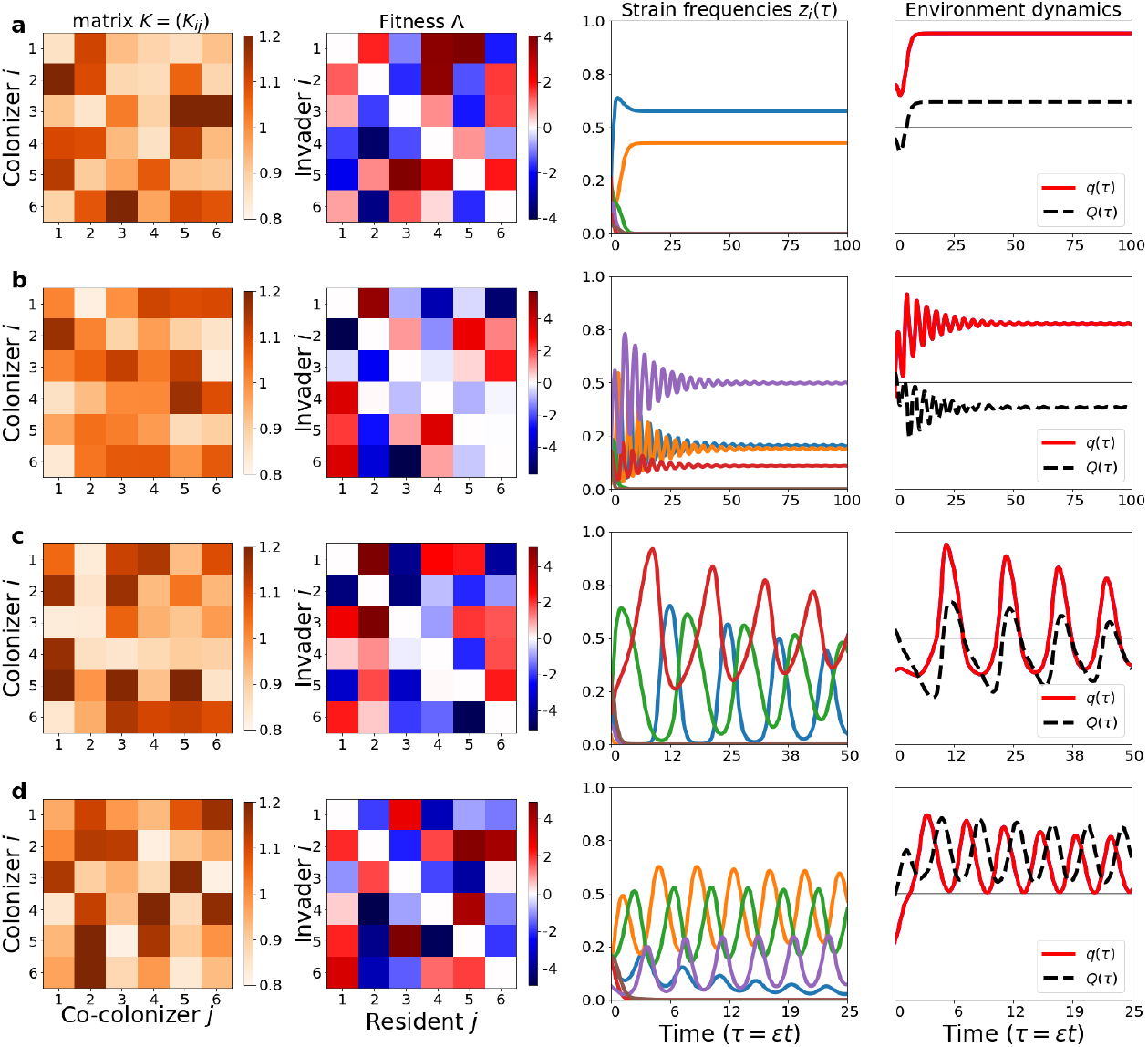
Evolutionary dynamics of mean traits in the *N*-strain system. **a.-b** Two cases of *R*_0_ = 3, where the dynamics of *q* and *Q* can be correlated or anti-correlated. Evolution of system resilience to invasion (*Q* high) can be obtained with evolution of more cooperation or more competition in the system, depending on initial conditions. **c-d.** Two cases of *R*_0_ − 2, where since *μ* is higher, the dynamics are more complex, and so is evolution of mean traits in the system. Remarkably *q* and *Q* can be synchronous or asynchronous. This regime corresponds more likely to maintenance of polymorphism in interaction trait space within the same population.

### S2 Mathematical details for the limits of *μ* → 0 and *μ* → ∞

We use the following notations, for the basic reproduction number and for the global steady state obtained via the neutral system:

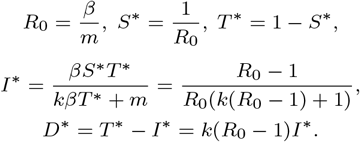

We define the important ratio *μ* of single and dual colonization by

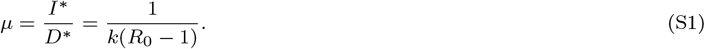

The reduced system reads

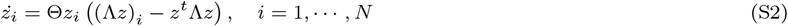

with the constraint

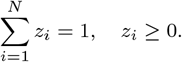

The *qualitative* behavior depends on the fitness *N* × *N* matrix 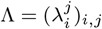 whose *i*,*j* term is the fitness of strain *i* with respect to the strain *j* and is explicitly given by

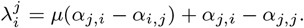

Here, *A* = (*α_ij_*) = ((*K_ij_* − *k*)/*ε*) is the normalized interaction matrix, accounting for deviation from neutrality.

Recall that 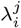 denotes the growth rate of the strain *i* when introduced to the steady state of the strain *j* alone: *E_j_* = (*z*_1_,⋯, *z_N_*) where *z_j_* = 1, *z_k_* = 0 for *k* ≠ *j*.

The time scale of the selective dynamics between multiple strains, in this system, depends basically on the constant

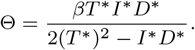

Our goal here is to give a description of the system in the limit cases *μ* → 0 and *μ* → +∞.

There are 3 parts in the discussion.

i. Which parameters change? *μ* depends on *k*, *β* and *m*. This point is straightforward from the formula (S1).
ii. The qualitative behaviour. This depends only on Λ (up to a positive multiplicative constant) which depends only on *μ*. Hence the point (*i*) has no importance here.
iii. How does the speed of the dynamics (Θ||Λ||) vary? This speed is especially important when it is close to 0. Remember that this system has been obtained by focusing of the first order perturbation in the quasi-neutral approximation (the zero order perturbation, *ε* = 0, yields exactly the neutral model). If the speed of the dynamics is below some threshold *O*(*ε*), then we cannot remove the next order perturbation and our model does not describe well the dynamics. In other words, if the speed is too slow then the system will stay very close to neutrality for a very long time. We will say that the system is *effectively neutral*. Hence, the main question in this section is the following: *May the speed of the dynamics go to zero when *μ* changes? If yes, what is the relevance of the parameters in point (i)?*

#### S2.1 Limit 1: Co-colonization, *μ* → 0: *k* → +∞ or *R*_0_ → +∞

##### The qualitative dynamics

Recall that 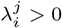 means that the strain *i* (the invader) may invade the system from a very small concentration within a system where only the strain *j* (the resident) is present.

Passing to the limit *μ* → 0, we obtain the matrix Λ_lim_ with the (*i, j*) coefficient being simply

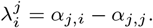

In general, very little may be said, because the structure of the matrix Λ_lim_ can be anything. As an example, in the special case *α_jj_* = 0 for all *j*, then Λ_lim_ = *A^T^*. Since *A* may have any structure, so can Λ_lim_, and very few results are known for the general replicator system. However, it is useful to see *A* as a random matrix and to check the probability of a given type of dynamics (see Yoshino et al. (2008)). We have the following interpretation.

- The fitness 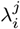 depends only on the coefficients *α_j,i_* and *α_j,j_* which are characteristics of the resident. *α_j,i_* measures how much the resident *j* contributes to the pool of co-colonization by the strains *i* and *j*: *D_ij_*, whereas *α_j,j_* measures how much the resident *j* increases the pool of self co-colonization: *D_jj_*.
- Hence, the relative fitness of the invader *i* depends only of the resident *j*. The invader *i* may invade the resident *j* if and only if the resident reinforces more the co-colonization *D_ij_* than *D_jj_*.
- This phenomenon provides a lot of niches for multiple strains to coexist. As an example, if *A* has a strong diagonal: *α_j,j_* > *α_i,j_* for any *j* = 1, ⋯ *N* and *i* ≠ *j* then all the strains may be stable residents when alone, and we will have at least *N* stable monomorphic steady states.
- In conclusion, let *A* be chosen randomly. The system is likely to have a lot of stable steady states wherein only few strains coexist. It is rare to see cycles or more complex dynamics.

##### The speed of the dynamics

Since *μ* → 0 then *μ* is bounded and far from 0, and so is ||Λ||. The speed of the dynamics is then measured by

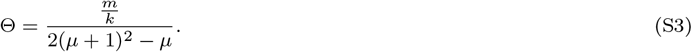

- If *k* → +∞, from the second expression in (S3) we have: 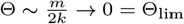. This is a problematic situation. *If the second infection occurs very quickly then* Θ *is very small and the model does not capture the real dynamics*.
- If *m* → 0 then again the same expression give 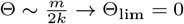. *If the natural mortality or the clearance of infection happen very slowly (which means that the rate of population turnover is very slow) then this model do not capture the real dynamics*.
- If *β* → +∞ then the same equation above shows that Θ → *m*2*k* = Θ_iim_ > 0 which, in this case, is far from zero. *Thus, if the pathogen transmission rate is very large, then the system captures well the real dynamics and the qualitative study of μ* → 0 *is appropriate*.

In summary, the only way for Θ_lim_ to be positive as *μ* → 0 is when *β* → +∞.

#### S2.2 Limit 2: single colonization, *μ* → +∞: *k* → 0 or *R*_0_ → 1

In that cases, we have generically ||Λ|| → +∞ and Θ → 0. In order to keep a bounded matrix Λ we rewrite the system as

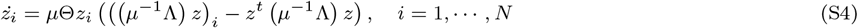

The speed is then given by *μθ* and the qualitative behavior by *μ*^−1^ Λ.

##### The qualitative dynamics

When *μ* → +∞, the matrix *μ*^−1^ Λ → Λ_lim_ = *A^T^* − *A* This matrix of pairwise invasion fitnesses becomes skew symmetric.

As opposed to the previous limit, there exist some results for the skew symmetric case. In the particular case where the coefficients are ±1 the system is known as the *Tournament game* which has been studied in detail in in Fisher and Reeves (1995) and has been used recently in Allesina and Levine (2011) in coexistence theory. Some of these results remain true for a general skew matrix Chawanya and Tokita (2002).

- The quadratic term in the frequency evolution equation is always^1^ zero. So the replicator equation system for *N* strains reduces to

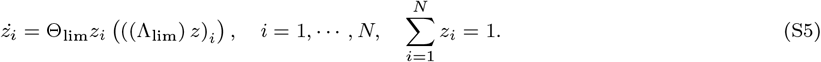
- Generically, there exists exactly one non-negative linearly stable equilibrium (in particular multistability is impossible). Without loss of generality, we can assume that this equilibrium reads *ζ* = (*ζ*_1_,⋯, *ζ_n_*, 0,⋯, 0) with 1 ≤ *n* ≤ *N*.
- From the linear point of view, *ζ* is a center: it is linearly stable but not asymptotically linearly stable. Indeed, the *N* − *n* last eigenvalues are negative and the *n* first eigenvalues are purely imaginary.
- The function 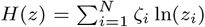 is bounded and increasing. We have 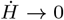 and 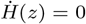 if and only *z_k_* = 0 for each *k* > *n*.
- It follows that *z_k_* → 0 for each *k* > *n*, and that the sub system of the *n* first strains is strongly persistent.
- In practice, exactly like the classical Lotka-Volterra model of prey and predators, we have a one-parameter family of cycles, parametrized by initial conditions, more precisely by the value of *H*(*z*(0)).
- Denote 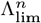 the sub-matrix of the first *n* rows and *n* columns of Λ^∞^ and *ζ_n_* = (*ζ*_1_,⋯, *ζ_n_*). We have

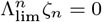 In particular it is necessary that 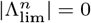. Since 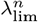 is skew symmetric, we have

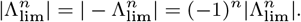 Hence, if *n* is odd we always have 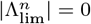. On the contrary, if *n* is even 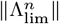 has no reason to be zero (except in very specific cases which have a zero probability of occurrence in the sense below) and there will be no steady state of *n* coexisting strains.
- A more complete description may be given. Let the coefficient of Λ_lim_ be chosen randomly. Denote *n* the number of strains coexisting, then the probability to have a steady state of *n* = *k* strains over a total pool of *N* strains is

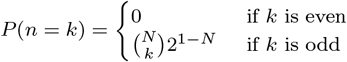 We illustrate this case in the figure S1. We finish this section by highlighting the fact that the above-described dynamics is not structurally stable. This structure will be lost for a large but finite value of *μ*. As *μ* decreases from +∞, the center will become either a stable focus or an unstable focus. In the first case, we will observe a global attractor, while in the second case, which is the most probable as shown in Figure 5, we will observe more complex dynamics like heteroclinic cycle or even chaos.

##### The speed of the dynamics

As shown below, the speed, in this second limit, is given by

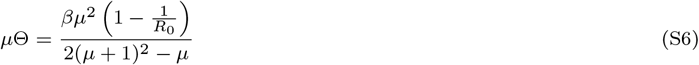

- If *R*_0_ → 1 then^2^ *μ*Θ → 0 = Θ_lim_ and the model is effectively neutral. *If the transmission intensity of the infection is too low, then this model do not capture well the dynamics*.
- If *k* → 0 then 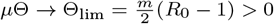 which is far from zero.

In summary: the only way for Θ_lim_ to be positive as *μ* → +∞ is to have *k* → 0, thus in the limit of extreme competition between strains in co-colonization. In that case, the compartment of co-colonization *D* becomes very small, and the dynamics are well described by the approximation *μ* → +∞, which depends only on the asymmetries *α_j,i_* − *α_i,j_*.

## Footnotes

1 The quadratic part is the scalar *Q*(*z*) = *z^T^* Λ*z*. From Λ = −Λ we obtain *Q*(*z*) = *Q*(*z*)*T* = *z^T^* Λ^*T*^ *z* = −*Q*(*z*), thus *Q*(*z*) = 0.

2 Except if *β* → ±∞ and 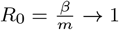, like by example if *β* = *m* + *a*. This corresponds to a situation where both the mortality-clearance rate and the infection rate become simultaneously very large. In that case, we rewrite 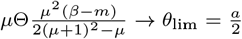 which is positive.

